# Programmable Editing of Primary MicroRNA Switches Stem Cell Differentiation and Improves Tissue Regeneration

**DOI:** 10.1101/2024.07.19.604260

**Authors:** Vu Anh Truong, Yu-Han Chang, Yi Tu, Thi Kieu Nuong Nguyen, Ngoc Nam Pham, Chin-Wei Chang, Yi-Hao Chang, Thi Kim Dung Ngo, Huu Dang Pham, Jui Tu, Thuc Quyen Dang, Anh Vy Truong, Yu-Chen Hu

**Author notes:** These authors contributed equally to this work (Vu Anh Truong, Yu-Han Chang).

## Abstract

Programmable RNA editing is harnessed for modifying mRNA. Besides mRNA, miRNA also regulates numerous biological activities, but current RNA editors have yet to be exploited for miRNA manipulation. To engineer primary miRNA (pri-miRNA), the miRNA precursor, we present a customizable editor REPRESS (RNA Editing of Pri-miRNA for Efficient Suppression of miRNA) and characterize critical parameters. The optimized REPRESS is distinct from other mRNA editing tools in design rationale, hence enabling editing of pri-miRNAs that are not editable by other RNA editing systems. We edited various pri-miRNAs in different cells including adipose-derived stem cells (ASCs), hence attenuating mature miRNA levels without disturbing host gene expression. We further developed an improved REPRESS (iREPRESS) that enhances and prolongs pri-miR-21 editing for at least 10 days, with minimal perturbation of transcriptome and miRNAome. iREPRESS reprograms ASCs differentiation, promotes in vitro cartilage formation and augments calvarial bone regeneration in rats, thus implicating its potentials for engineering miRNA for many applications such as stem cell engineering and tissue regeneration.

## Introduction

Programmable RNA editing enables transient gene regulation without permanent genome change^1^ and promises a safer option than DNA editing^2^. Adenosine deaminase acting on RNA 2 (ADAR2) naturally deaminates adenine (A) to inosine (I) in double-stranded RNA (dsRNA)^3^ and preferentially deaminates a target A mismatched with a cytosine (C)^4^. For programmable RNA editing, deaminase domain of ADAR2 (ADAR2_DD_) has been fused to deactivated Cas13 (dCas13) protein, which is guided by a single CRISPR RNA (crRNA) comprising a spacer sequence that base pairs with the RNA substrate^5^.

The first system exploiting this strategy, REPAIR, comprises a fusion protein of dCas13b derived from *Prevotella sp. P5-125* (dPspCas13b) and hyperactive ADAR2_DD_. Coupled with a site-specific crRNA, REPAIR confers precise A->I conversion^6^. Subsequently, LEAPER employs engineered ADAR-recruiting RNAs (arRNAs) to recruit endogenous ADARs for A->I conversion^7^. RESTORE exploits synthetic antisense oligonucleotides to recruit endogenous ADARs^8^. Recent studies also exploit short, chemically modified oligonucleotides or circular RNA to recruit endogenous ADAR^9, 10^. Other programmable RNA-editing tools such as REPAIRx^11^, CURE^12^, REWIRE^1^, RESCUE^4^ and CIRTS^13^ were also developed. Site-specific RNA editing can also be achieved by fusing ADARs with an RNA-binding peptide or oligonucleotide^14^. Most of these tools edit mRNA by designing the guide RNA or oligonucleotide complementary to mRNA except a single mismatch (usually a C) opposite to the adenine intended for deamination. These methods are harnessed to reverse pathogenic mutations on mRNA, restore native protein expression in cells and disease models. However, they are neither used to edit primary microRNA (pri-miRNA), nor have they been exploited for tissue regeneration.

Pri-miRNA is the miRNA precursor co-expressed with its host genes in the nucleus. Pri-miRNA folds up into an imperfect hairpin containing a double-stranded stem-loop region^15^. Nascent pri-miRNA is cleaved near the junction (called basal junction) of single stranded RNA (ssRNA) and dsRNA hairpin by Microprocessor composed of Drosha/DGCR8 into a precursor miRNA called pre-miRNA^16^. Once processed, pre-miRNA is exported out of nucleus and undergoes further maturation to form the mature miRNA. miRNA functions as the guide molecule in RNA silencing to regulate developmental and pathological processes.

Because pri-miRNA processing is critical for miRNA biogenesis and current programmable RNA editing systems have yet to be harnessed to edit pri-miRNA, this study aims to create a programmable system for <u>R</u>NA <u>E</u>diting of <u>PR</u>i-miRNA for <u>E</u>fficient <u>S</u>uppre<u>S</u>sion of miRNA (REPRESS). The optimized REPRESS enables editing of various pri-miRNAs and attenuates their mature miRNAs levels in different cells including adipose-derived stem cells (ASCs). We further generate the improved REPRESS (iREPRESS) that prolongs and enhances pri-miRNA editing with minimal perturbation of transcriptome and microRNAome. iREPRESS editing of pri-miR-21 reprograms ASCs differentiation, ameliorates *in vitro* cartilage formation and *in vivo* bone regeneration, thus paving a new avenue for miRNA regulation, stem cell engineering and regenerative medicine.

## Results

### Development of pri-miRNA base editor (REPRESS)

To edit pri-miRNA in a programmable fashion, we created REPRESS by fusing dCas13 and human ADAR2 deaminase domain (ADAR2_DD_) with a linker, in combination with a crRNA to guide the editor to the pri-miRNA (Fig. 1A). Unlike other ADAR-based RNA editing methods that introduce an A-C mismatch at the crRNA:mRNA duplex^6–8^, we designed the crRNA spacer sequence perfectly complementary to ssRNA sequences near the basal junction of pre-miRNA hairpin, such that ADAR2_DD_ deaminates adjacent pre-miRNA hairpin (Fig. 1A).

**Fig. 1:**
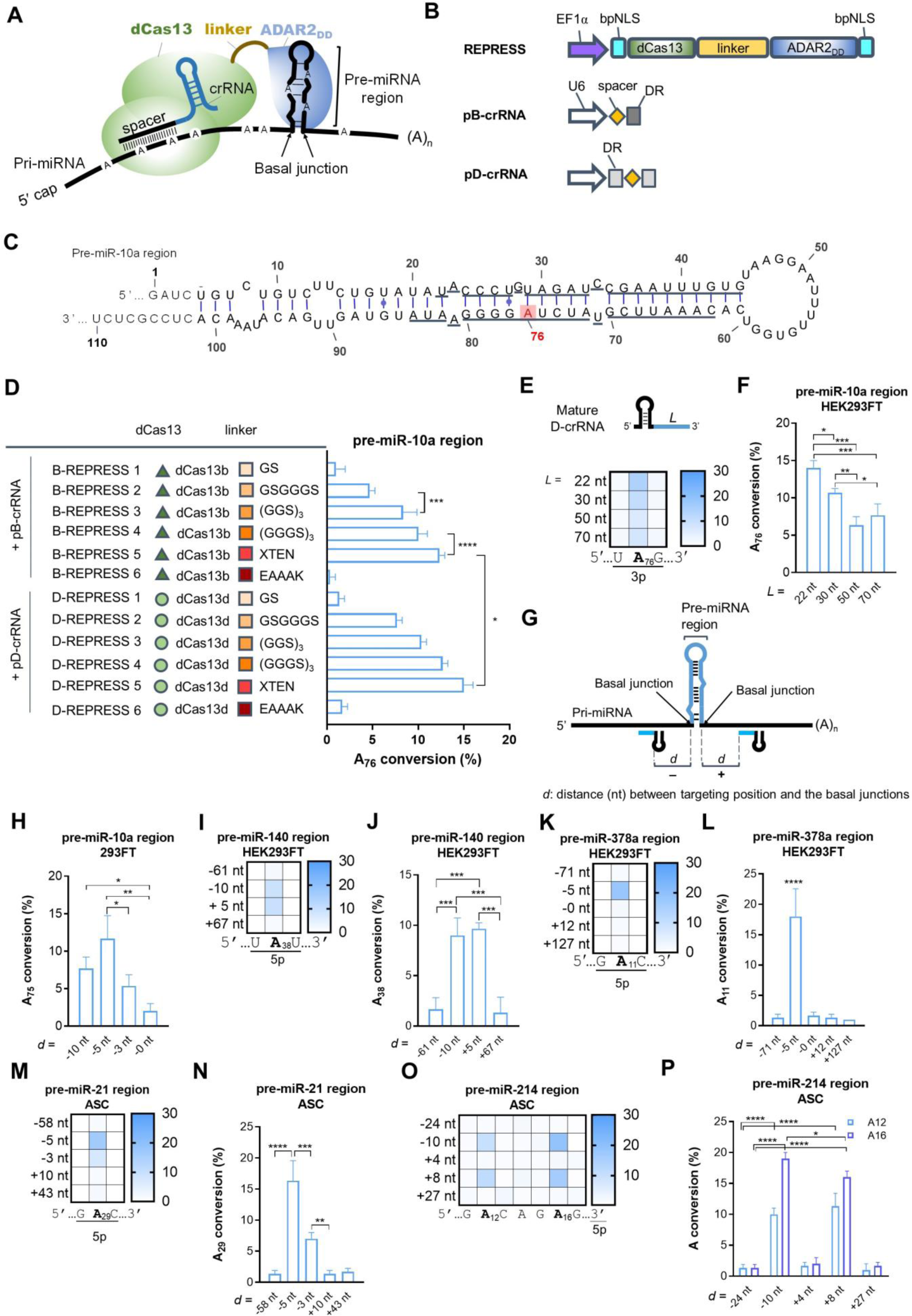
Development of REPRESS for pri-miRNA editing. (A) Mode of action of REPRESS. crRNA targeting to the ssRNA motif of pri-miRNA brings the dCas13-ADAR2_DD_ to the vicinity of basal junction, thus catalyzing specific A->I conversion within the pre-miRNA hairpin. (B) The REPRESS expression cassette. dCas13 (dPspCas13b or dRfxCas13d) was fused with ADAR2_DD_ via various linkers. The fusion protein was flanked by two bipartite nuclear localization signals (bpNLS) to facilitate nuclear import. crRNA expression cassette was driven by U6 promoter with an architecture of direct repeat (DR)-spacer for dPspCas13b (pB-crRNA) or DR-spacer-DR for dRfxCas13d (pD-crRNA). (C) Secondary structure of pre-miR-10 hairpin. The numbering of bases was defined according to the sequences retrieved from miRBase database. (D) REPRESS variants encoding dPspCas13b (B-REPRESS 1∼6) or dRfxCas13d (D-REPRESS 1∼6) and various linkers. Corresponding editing efficiencies are also shown. (E, F) Editing efficiency using different spacer lengths. (G) Illustration of relative distance (*d*) between targeting position and the 5’ (‒) or 3’ (+) basal junctions. (H-P) Editing efficiencies at different targeting positions for various pri-miRNAs. Plasmids encoding REPRESS and crRNA were co-delivered into cells with a ratio of 1:3 (w:w). Total RNA was harvested after 1 day for reverse transcription and PCR amplification. PCR amplicons encompassing the pre-miRNA and flanking sequences were Sanger sequenced. A->I editing efficiencies were calculated by EditR. Data represent means±SD of three independent culture experiments. ****p < 0.0001; ***p < 0.001; **p<0.01; *p<0.05.

For proof-of concept, we first generated a series of REPRESS plasmids by fusing dPspCas13b or dRfxCas13d with hyperactive ADAR2_DD_ using different linkers to edit pri-miR-10a abundantly expressed in HEK293FT cells (Fig. 1B-C). The fusion gene was flanked with bipartite nuclear localization signals (bpNLS) to ensure editing of nuclear pri-miRNA. The crRNA plasmids associated with dPspCas13b (pB-crRNA) or dRfxCas13d (pD-crRNA) were constructed to guide REPRESS to the proximity of pre-miR-10a basal junction. One day after plasmid co-transfection, the A->I editing events were measured by Sanger sequencing of RT-PCR amplicons (444 bp) using primers flanking the pre-miRNA hairpin (Extended Data Fig. 1).

When we used dPspCas13b and the GS linker (B-REPRESS 1, Fig. 1D, Extended Data Fig. 1), in the entire amplicon we only detected marginal A->I conversion at A at position 76 (A_76_) in the pre-miRNA region (Fig. 1C-D). No other editing events were found elsewhere in the amplicon. By increasing the lengths of GS-based linker to 6-aa (GSGGGS), 9-aa (3×GGS) and 12-aa (3×GGGS), we detected elevating A->I conversion efficiency, but only at A_76_ among 28 adenines in the pre-miR-10a hairpin (Fig. 1D and Extended Data Fig. 1). The 16-aa flexible XTEN linker (B-REPRESS 5) further improved A_76_ conversion efficiency to ≍12%. However, the 5-aa rigid linker (EAAAK) barely converted A_76_.

When we replaced dPspCas13b with dRfxCas13d (D-REPRESS 1∼6, Fig. 1D), the D-REPRESS editors conferred A_76_ conversion in a similar trend, but with generally higher efficiency when using the same linker (Fig. 1D and Extended Data Fig. 2). Regardless of dCas13 type and linkers, only A_76_ was converted (Extended Data Fig. 1-2). Among these editors, D-REPRESS 5 that exploited XTEN linker to fuse dRfxCas13d with ADAR2_DD_ conferred the highest A_76_ editing efficiency (≍15%). Therefore, we abbreviated D-REPRESS 5 as REPRESS for ensuing characterization.

dRfxCas13d crRNA consists of a stem-loop and a spacer sequence for base pairing the target RNA. Because transfection of ≍70 nt synthetic oligonucleotide to base pair mRNA can recruit endogenous ADAR for mRNA editing^7^, we were inspired to vary the crRNA spacer length (L, Fig. 1E). We found that increasing the spacer length from 22 to 70 nt all enabled A_76_ conversion, but with descending efficiencies (Fig. 1E-F), indicating that 22 nt is the most optimal spacer length. We next tested whether changing the crRNA targeting position would alter the A->I conversion preference. We fabricated two crRNA spacers, one complementary to pre-miR-10a with an A-C mismatch at A_76_, the other with the highest predicted dRfxCas13d targeting score. Both designs, however, did not achieve A->I conversion (Extended Data Fig. 3). Other RNA editing systems such as REPAIR, LEAPER and RESTORE that introduced an A-C mismatch targeting A_76_ (Extended Data Fig. 3) or targeted the ssRNA adjacent to the 5’ basal junction of pre-miR-10a also failed to achieve A->I conversion (Extended Data Fig. 4).

We further designed crRNAs to target bases with different distances between the crRNA binding site and the basal junction (*d*, Fig. 1G). We uncovered that *d* plays crucial roles to dictate the conversion rate although still only A_76_ was converted. crRNA targeting the basal junction (*d*=0) conferred the poorest A_76_ conversion while increasing the distance to 5 nt (*d*=−5) yielded the highest A_76_ conversion rate (≍12%, Fig. 1H). In HEK293FT cells, REPRESS also enabled A->I conversion of pre-miR-140 at A_38_ (Extended Data Fig. 5) with the highest editing efficiency at *d*=+5 or −10 (Fig. 1I-J). Conversely, REPRESS converted A_11_ of pre-miR-378a with the highest efficiency reaching ≍18% at *d*=−5 (Fig. 1K-L, Extended Data Fig. 5). We further detected A_29_ conversion for pre-miR-21 (Extended Data Fig. 5) abundant in adipose-derived stem cells (ASCs), whose highest efficiency reached ≍17% at *d*=−5 (Fig. 1M-N). By contrast, none of the REPAIR, LEAPER and RESTORE systems achieved A_29_ conversion, regardless of whether the guide RNAs targeted the pri-miR-21 hairpin or ssRNA sequences adjacent to the basal junction (Extended Data Fig. 6-7). For pri-miR-214 in ASCs, we detected conversion at two adenines (A_12_ and A_16_), with the highest efficiency approaching 19% at *d*=−10 (Fig. 1O-P). These two adenines are in the bulge, which can be deaminated by ADAR^17^.

### Editing of pri-miRNAs by REPRESS suppressed mature miRNAs levels

We next evaluated whether REPRESS-directed pri-miRNA editing inhibited miRNA maturation, using TaqMan qPCR primers specific to mature miRNA. As a control, we constructed a deactivated REPRESS (dREPRESS) by fusing dRfxCas13d with deactivated ADAR2_DD_ (Fig. 2A). Using the optimal crRNA targeting design (Fig. 1), dREPRESS partially reduced the levels of several miRNAs in different cells (Fig. 2B-G). REPRESS-mediated A_76_ conversion at pre-miR-10a (3p strand) further inhibited miR-10a-3p level for 56% (Fig. 2B). A_38_ conversion at pre-miR-140 (5p strand) and A_11_ conversion at pre-miR-378a (5p strand) significantly knocked down miR-140-5p and miR-378a-5p levels for 25% and 79%, respectively (Fig. 2C-D). We also targeted the same pri-miR-10a region in A549 lung cancer cells and achieved ≍60% inhibition of miR-10a-3p level (Fig. 2E). The editing inhibited A549 cell proliferation, migration and invasion (Extended data Fig. 8). In ASCs, A_29_ and A_12_/A_16_ conversion at pre-miR-21 (5p strand) and pre-miR-214 (5p strand) suppressed miR-21-5p and miR-214-5p levels for 72% and 58%, respectively (Fig. 2F-G). Furthermore, pri-miRNAs are usually co-expressed with their host genes. Editing of pri-miR-10a (in HEK293FT) and pri-miR-21 (in ASCs) did not disturb the levels of their host genes (Extended Data Fig. 9).

**Fig. 2:**
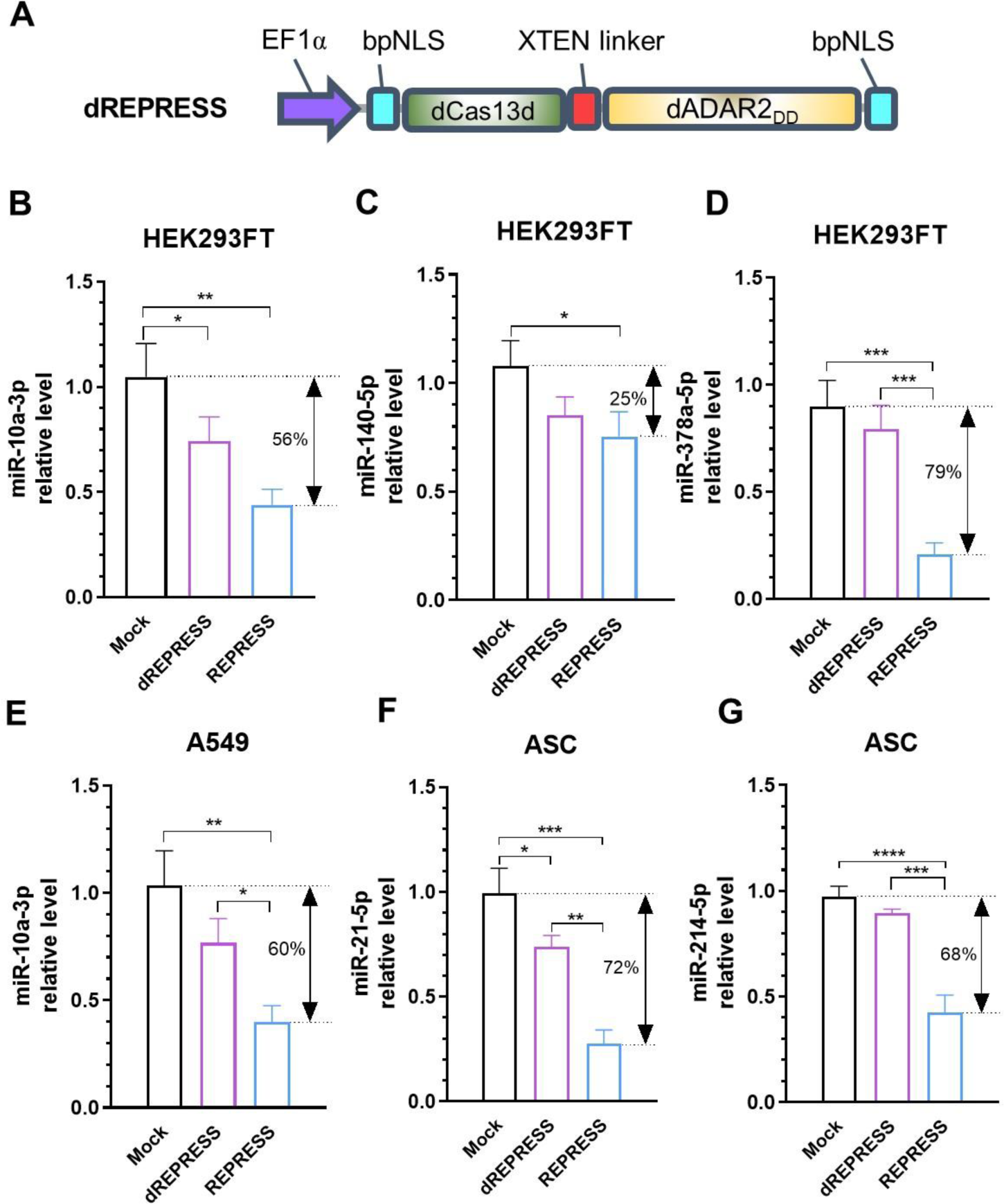
Editing of pri-miRNAs knocked down various mature miRNAs in different cells. (A) dREPRESS expression cassette harboring the fusion of dRfxCas13d and deactivated ADAR2_DD_ with E488Q and E396A mutations. (B-G) Relative mature miRNAs levels. Plasmids encoding optimized REPRESS or dREPRESS were co-delivered with crRNA expressing plasmids into cells with a ratio of 1:3 (w:w). miRNAs were collected at 1 day for TaqMan assay using primers specific to mature miRNAs. Data represent means±SD of three independent culture experiments. ****p < 0.0001; ***p < 0.001; **p<0.01; *p<0.05.

### Improved REPRESS (iREPRESS) for prolonged pri-miRNA editing and safety profiling

We reasoned that prolonged pri-miRNA editing may extend future therapeutic efficacy. To this end, we packaged REPRESS into a hybrid Cre/loxP-based baculovirus (BV) system that transduces ASCs at >95% efficiency and prolongs transgene expression^18^. The hybrid vector comprises two BV: one expressing Cre recombinase and the other harboring the transgene flanked by loxP sites^18^. Co-transduction of ASCs with the two BV enables intracellular loxP-flanking transgene re-circularization and prolongs transgene expression^18^.

We generated BV vectors Bac-REPRESS encoding REPRESS and Bac-cr21 expressing cr21 to target pri-miR-21 at *d*=−5 as in Fig. 1F (Fig. 3A). Both REPRESS and cr21 cassettes were flanked by two *in cis* loxP sequences. We co-transduced ASCs with Bac-REPRESS and Bac-cr21 (REPRESS group) and detected 12% A_29_ conversion at 3 days post-transduction (dpt), which rapidly decreased with time (Fig. 3B). When we co-transduced ASCs with Bac-REPRESS/Bac-cr21 and a third Bac-Cre (Fig. 3A) that expressed Cre (iREPRESS group), A_29_ conversion rate increased to ≍23% at 3 dpt and remained 16% at 7 dpt and 5% at 10 dpt, indicating that iREPRESS enhances and prolongs A_29_ conversion (Fig. 3B). Concurrently, iREPRESS more effectively knocked down miR-21-5p levels for at least 10 days (Fig. 3C).

**Fig. 3.**
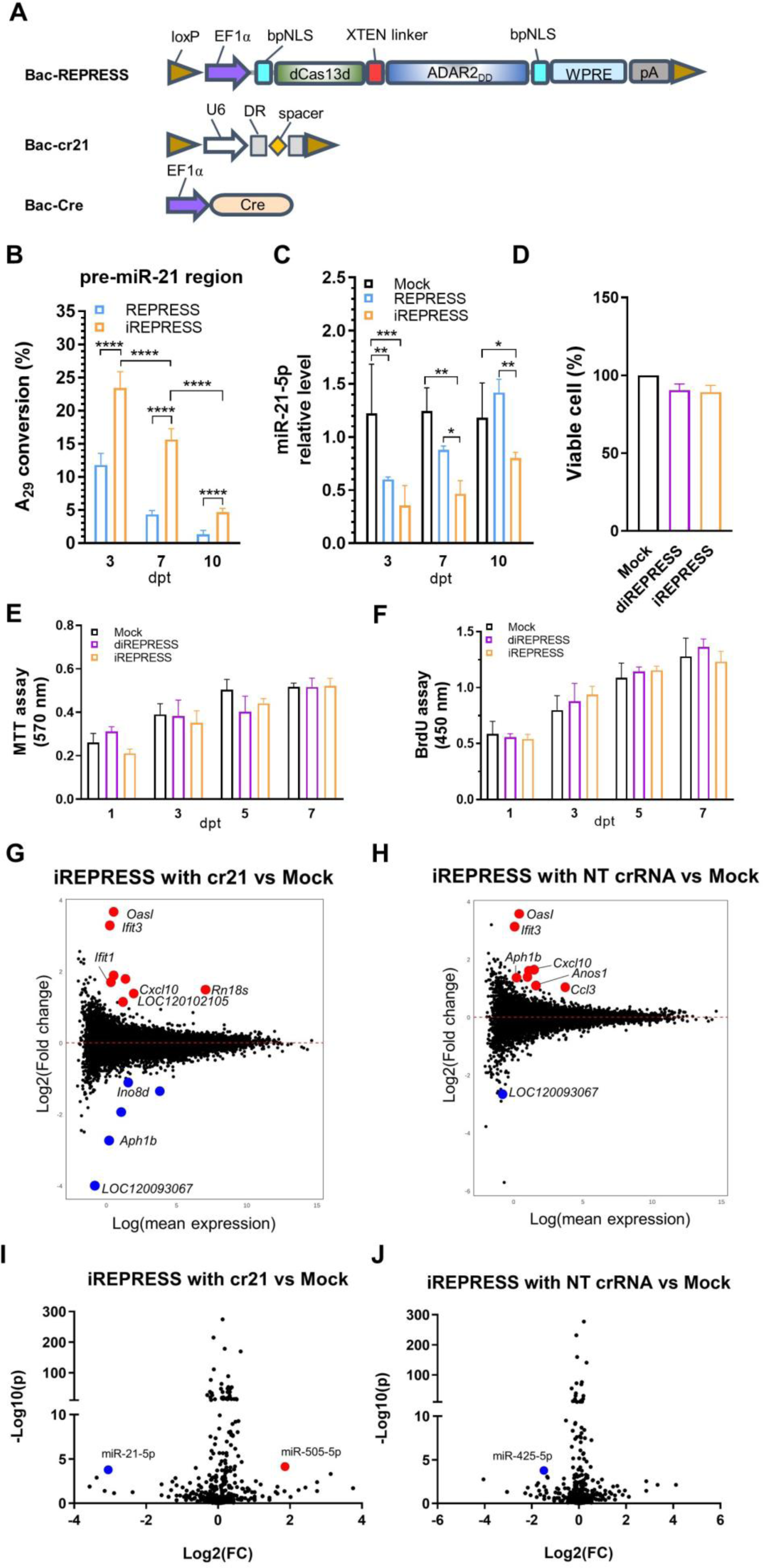
Development of improved REPRESS (iREPRESS) and safety profiling. (A) Illustration of BV vectors. Bac-REPRESS expressed the entire REPRESS cassette flanked by two loxP sequences. Bac-cr21 expressed the crRNA targeting pri-miR-21 at *d*=−5 near the 5′ basal junction of the pre-miR-21 hairpin. Bac-Cre expresses Cre recombinase to excise and circularize the loxP-flanked REPRESS for sustained expression. ASCs were Mock-transduced (Mock group), co-transduced with Bac-REPRESS/Bac-cr21 (REPRESS group) or co-transduced with Bac-REPRESS/Bac-cr21/Bac-Cre (iREPRESS group). (B) A_29_ conversion rate as measured by deep sequencing. (C) Relative mature miR-21-5p level measured by TaqMan assay. (D) Viable cell number. (E) MTT assay. (F) BrdU incorporation assay. (G-H) Transcriptome-wide analysis of ASCs co-transduced with iREPRESS using cr21 (G) or non-targeting (NT) crRNA (H), with the Mock group as the reference. RNA-seq data are presented as log2(fold change) vs. log(mean expression). (I-J) Global miRNA analysis of ASCs co-transduced with iREPRESS using cr21 (I) or NT crRNA (J). Small RNA-seq data are presented in ‒log10(*p* value) vs. log2(fold change). Data represent means±SD of three independent culture experiments. ****p < 0.0001; ***p < 0.001; **p<0.01; *p<0.05; Red and blue dots represent significantly upregulated and downregulated genes in sequencing data.

To evaluate the safety profile, ASCs in the iREPRESS group were analyzed and referenced to the Mock group. As a control, dREPRESS (Fig. 2A) was packaged into BV for co-transduction with Bac-cr21/Bac-Cre (diREPRESS group). Both diREPRESS and iREPRESS groups did not mitigate ASCs viability (Fig. 3D). Neither did they compromise cell metabolism and proliferation (Fig. 3E-F). We next performed transcriptome-wide RNA-seq to assess off-target effects of iREPRESS in rat ASCs. Compared with the Mock group, iREPRESS triggered minimal transcriptome perturbations (*p*<0.05) with only 8 upregulated and 5 downregulated genes (Fig. 3G). When we replaced cr21 with a non-targeting crRNA (NT crRNA), only 9 genes were perturbed (Fig. 3H). Small RNA sequencing showed that iREPRESS specifically inhibited miR-21-5p levels with only one off-target miR-505-5p upregulation (Fig. 3I). Replacing cr21 with NT crRNA abolished miR-21-5p downregulation, with only one off-target downregulation of miR-425-3p (Fig. 3J). These data confirmed that iREPRESS prolonged pri-miRNA editing without evident perturbation of transcriptome and miRNAome.

### Switching ASCs differentiation by iREPRESS-prolonged pri-miR-21 editing

ASCs can differentiate towards adipogenic, chondrogenic and osteogenic lineages, making them useful for cartilage^19^ and bone^20^ regeneration. However, ASCs differentiate favorably towards adipogenic lineage, partly because endogenous miR-21 promotes adipogenesis, hence mitigating ASCs’ potential for cartilage/bone regeneration^21^. Therefore, we next exploited iREPRESS to reprogram ASCs differentiation.

ASCs were Mock-transduced (Mock group), co-transduced with Bac-REPRESS/Bac-cr21 (REPRESS group) or with Bac-REPRESS/Bac-cr21/Bac-Cre (iREPRESS group). iREPRESS enabled more significant suppression of adipogenic marker genes *C/ebpα* and *Ppar-γ* (Fig. 4A-B) while eliciting stronger expression of chondrogenic markers *Acan* and *Col2a1* (Fig. 4C-D). Oil Red O staining for oil droplet formation (for adipogenesis), Alcian blue staining for glycosaminoglycan (GAG, for chondrogenesis), Alizarin Red staining for mineralization (for osteogenesis) and ensuing quantitative analyses (Fig. 4E-J) confirmed that iREPRESS suppressed adipogenesis and triggered chondrogenesis more potently than the REPRESS and Mock groups but did not apparently affect osteogenesis.

**Fig. 4:**
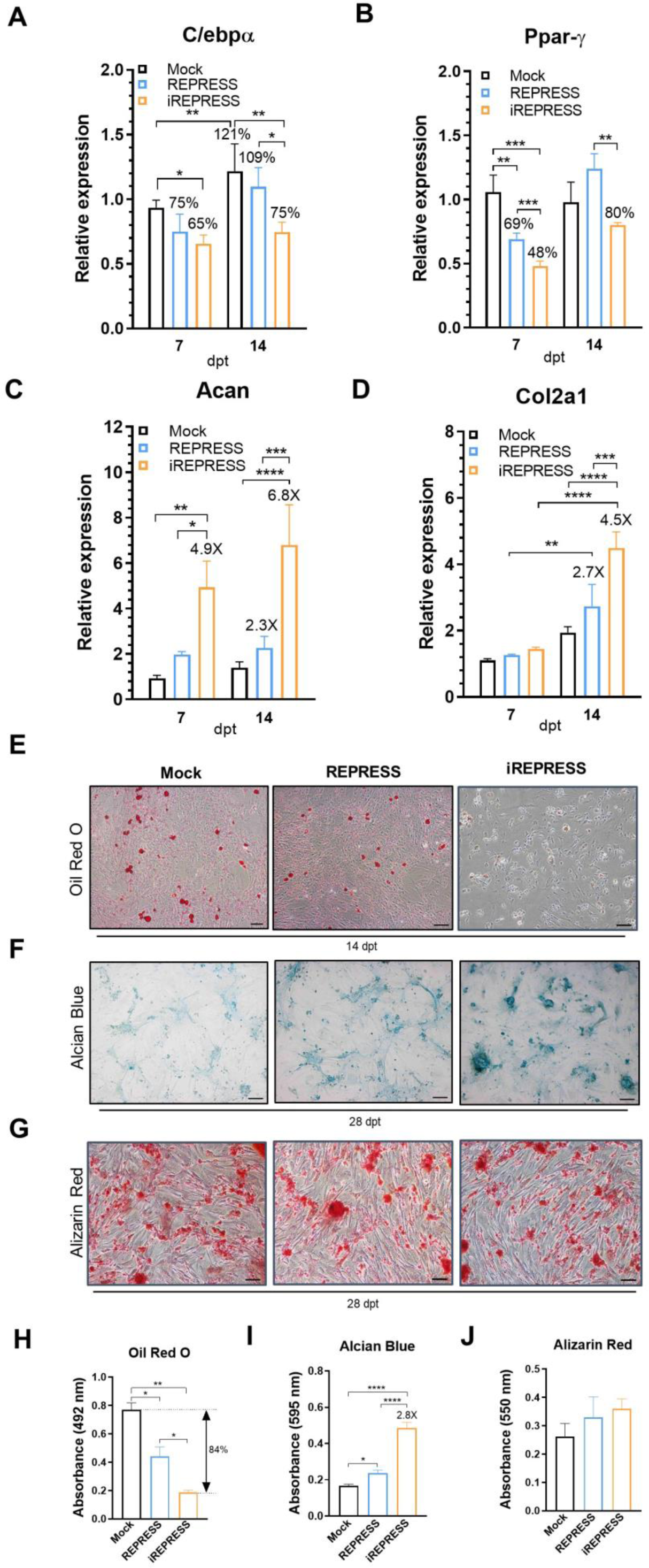
iREPRESS switched ASCs from adipogenic towards chondrogenic differentiation. Experiments and experimental groups are identical to those shown in Fig. 3. (A-B) Relative expression of adipogenic marker genes *C/ebpα* (A) and *Ppar-γ* (B). (C-D) Relative expression of chondrogenic marker genes *Acan* (C) and *Col2a1* (D). (E-G) Safranin O, Alcian Blue and Alizarin Red staining. (H-J) Spectrophotometric analysis for oil droplet formation (adipogenesis, H), GAG (for chondrogenesis, I) and mineralization (for osteogenesis, J). Data represent means±SD of three independent culture experiments. ****p < 0.0001; ***p < 0.001; **p<0.01; *p<0.05.

### iREPRESS-mediated pri-miR-21 editing in ASCs stimulated *in vitro* cartilage growth

Hyaline cartilage formation starts from stem cell condensation, chondrogenic differentiation and deposition of extracellular matrix (ECM) molecules such as collagen II (Col II) and GAG. We transduced ASCs as in Fig. 4, separately seeded the Mock and iREPRESS group cells into porous scaffolds (*n*=3 for each group) and cultured the ASCs/scaffold constructs. At 7 and 14 dpt, the iREPRESS group constructs exhibited whitish and glassy appearance that resembled the ECM of hyaline cartilage (Fig. 5A). H&E staining revealed the onset of condensation at 7 dpt and accumulation of more ECM in the iREPRESS group than the Mock group at 14 dpt (Fig. 5B). Alcian Blue and Col II immunostaining and quantitative analyses showed that iREPRESS deposited more uniform GAG and Col II than the Mock group at both 7 and 14 dpt (Fig. 5C-F).

**Fig. 5:**
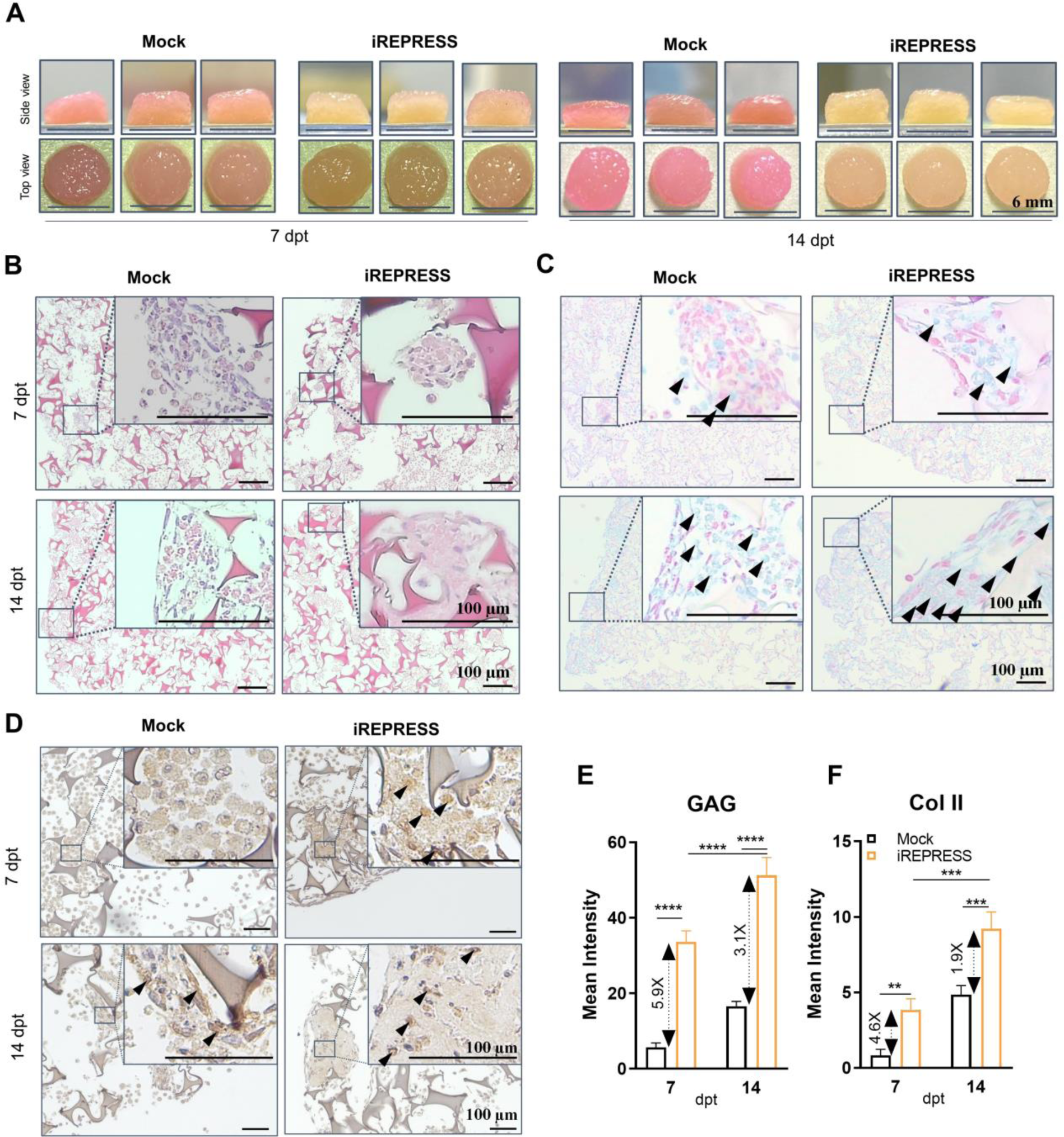
iREPRESS-mediated miR-21 knockdown enhanced cartilage formation. (A) Gross appearance of engineered cartilages. (B) H&E, (C) Alcian Blue and (D) Col II staining. (E-F) Semiquantitative analysis of GAG and Col II. ASCs were mock- (mock group) or iREPRESS-transduced (iREPRESS group) in 15-cm dishes. At 1 dpt, the cells were seeded into porous gelatin scaffolds (diameter=6 mm; thickness=1 mm; 5×10^6^ cells/scaffold; *n*=3 for each group). The ASCs/gelatin constructs continued to be cultured in chondroinductive medium and assayed at 7 or 14 dpt. Quantitative data represent means±SD of three independent culture experiments. ****p < 0.0001; ***p < 0.001; **p<0.01; *p<0.05.

### iREPRESS-mediated pri-miR-21 editing stimulated calvarial bone regeneration *in vivo*

Healing of large calvarial bone defect remains challenging for orthopedic surgeons. We previously showed that implantation of chondroinductive ASCs ameliorates calvarial bone healing by switching the repair pathway^22, 23^. To evaluate the feasibility of pri-miRNA-21 editing to promote calvarial bone regeneration, we harvested the Mock and iREPRESS constructs at 7 or 14 dpt (as in Fig. 5) and implanted them into critical-size calvarial defects in rats (*n*=4∼5 for Mock and *n*=6 for iREPRESS at both 7 and 14 dpt).

Micro CT (µCT) imaging and quantitative analysis revealed that the Mock constructs elicited slow and poor bone regeneration, regardless of harvesting at 7 or 14 dpt (Fig. 6), which underscores the difficulty to repair large calvarial defects using ASCs. The iREPRESS group, nonetheless induced formation of bone islands as early as week 4 (W4) and progressive formation of new bone (Fig. 6A-C). Quantitative data confirmed that the iREPRESS constructs elicits significantly faster and superior bone regeneration, with the nascent bone area, volume and density at W12 approaching 44-48%, 22-24% and 32-33% of the original defect, respectively (Fig. 6D-F). The *in vitro* culture time at which the constructs were collected (7 or 14 dpt) did not significantly influence the regeneration. Furthermore, histological and immune staining of the regenerated bone specimens showed evidently more bone formation, bone-specific ECM accumulation and less fibrous tissues in the iREPRESS group than the Mock group (Extended Data Fig. 10).

**Fig. 6:**
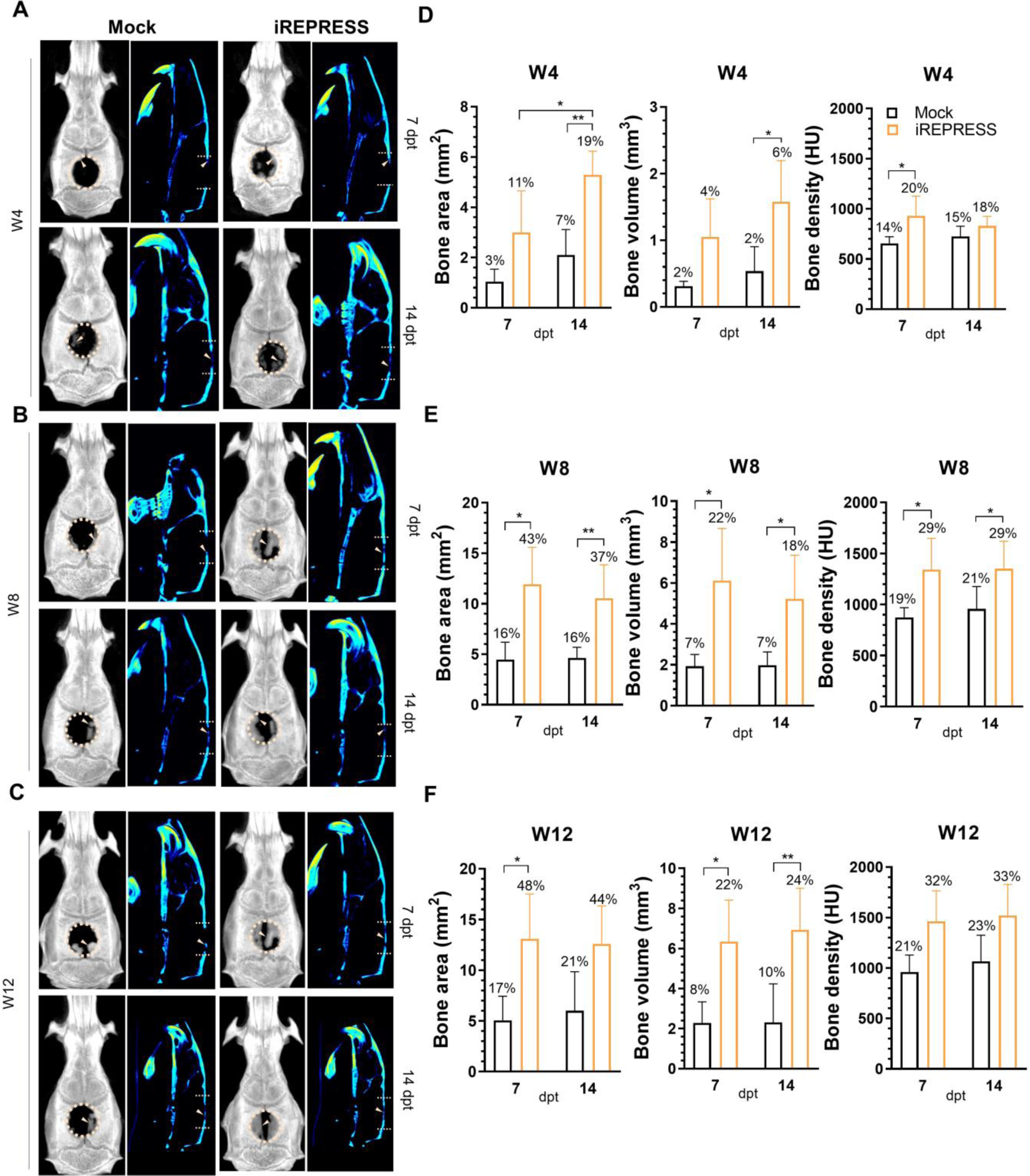
iREPRESS-engineered ASCs/scaffold constructs promoted calvarial bone healing. (A-C) Representative 3D and hot-and-cold projections of the transverse and sagittal views of the calvarial bone at 4 (A), 8 (B) and 12 (C) weeks post-implantation. (D-F) Regenerated bone area (mm^2^), volume (mm^3^) and density (HU, Hounsfield Unit). ASCs from the Mock and iREPRESS groups (as in Fig. 5) were seeded into gelatin scaffolds, cultured and harvested at 7 or 14 dpt. The constructs were implanted into the critical-sized (diameter=6 mm) calvarial defects of SD rats. Bone regeneration was evaluated by μCT imaging. Percentage values were calculated by normalization to the bone area (28 mm^2^), volume (28 mm^3^) and density (4600 HU) of the original defects. ****p < 0.0001; ***p < 0.001; **p<0.01; *p<0.05.

## Discussion

Current programable RNA editing tools are used for mRNA editing but have yet to be exploited to edit pri-miRNA. Most of these tools use a guide RNA carrying a A-C mismatch at the pre-determined A in mRNA to direct ADAR-mediated A->I conversion. We report the first programmable pri-miRNA base editor REPRESS. REPRESS distinguishes itself in that the crRNA spacer is completely complementary to the ssRNA near the basal junction of pre-miRNA hairpin, thus re-directing the dRfxCas13d-ADAR2_DD_ effector to deaminate adjacent pre-miRNA hairpin duplex (Fig. 1A-B). REPRESS is also distinct from other RNA editing tools in that REPRESS is designed to translocate to the nucleus where pri-miRNA is located, whereas other systems are introduced into the cytoplasm to edit mRNA.

We identified that REPRESS editing depends on dCas13 protein type, linker types and spacer lengths (Fig. 1D-F). dRfxCas13d generally confers higher editing efficiency than dPspCas13b used in the REPAIR system^6^, probably because Cas13d has a favorable accessibility toward RNA hairpin^24, 25^. Among all linkers tested, the 16-aa flexible XTEN yields the highest editing efficiency, which concurs with the notion that XTEN outperforms other linkers in DNA base editing^26^ and highlights the importance of linker length and flexibility for positioning ADAR2_DD_ to specific A in the pre-miRNA hairpin. Importantly, we identified the optimal crRNA targeting position in the window of 5 to 10 bases away from the pre-miRNA basal junction (Fig. 1G-P). These findings allow us to develop the optimal REPRESS that successfully edits various pri-miRNAs in different cells with efficiency varying with pri-miRNAs, presumably because ADAR editing site preference and efficiency hinge on different pri-miRNAs^3^. REPRESS preferentially edits A within the mature miRNAs, a phenomenon observed previously^3, 17^. Intriguingly, endogenous ADAR edits multiple adenines in pri-miRNAs^3, 17^ but REPRESS edits only one or two adenines. The discrepancy may arise because endogenous ADAR binds the dsRNA substrate and deaminates A depending on its own structure and interaction with dsRNA. In contrast, REPRESS is guided by crRNA to ssRNA motif adjacent to the basal junction of pre-miRNA hairpin, which may constrain the accessibility of ADAR2_DD_ active site to many A in the pre-miRNA hairpin. Further structural studies are required to decipher the rule of how REPRESS selects specific A for conversion. Nonetheless, the optimized REPRESS can edit different pri-miRNAs that are not editable by other RNA editing systems such as REPAIR, LEAPER and RESTORE (Extended Data Fig. 3-7), which may be attributed to the nuclear localization of REPRESS by bpNLS while other systems are located in the cytoplasm. These data underscore the stringent requirement for REPRESS design and indicate the superiority of REPRESS for pri-miRNA editing.

Despite editing at only one or two A, REPRESS-mediated editing replaces A-U Watson-Crick pair with mismatched I**·**U wobble pair, which causes substantial changes in the stem structure and hinders pri-miRNA cleavage by Drosha, hence inducing pri-miRNA degradation by a ribonuclease specific to inosine-containing dsRNAs^3, 17^. In accord, REPRESS-mediated editing knocks down different mature miRNAs in cell lines, cancer cells and stem cells (Fig. 2). The improved REPRESS (iREPRESS) prolongs and enhances pri-miRNA-21 editing and reduces mature miR-21 levels for 10 days. Notably, previous ADAR_DD_-associated RNA editing strategies usually suffer from evident off-target effects^11^, yet iREPRESS induces minimal off-target effects and perturbations on the transcriptome and microRNAome (Fig. 3), probably because iREPRESS is transported into nucleus, and nuclear localization of ADAR reduces off-target effects^27, 28^. Critically, miR-21 knockdown by iREPRESS reprograms the ASCs differentiation from adipogenic to chondrogenic pathway, promotes *in vitro* cartilage formation (Fig. 5) and stimulates calvarial bone healing (Fig. 6 and Extended Data Fig. 10).

RNA editing offers numerous opportunities for fundamental research and medical applications^29^. Biotechnology industry has begun to use RNA editing to develop treatments from genetic diseases to temporary maladies such as acute pain^2^. Here we demonstrate a new application of tissue regeneration by pri-miRNA editing using iREPRESS. miRNAs also regulate the expression of human genes responsible for numerous biological functions. Although miRNA sponges or other approaches are used to degrade miRNA, these methods may also induce host gene silencing. In contrast, iREPRESS specifically suppresses miRNA editing without affecting the host genes (Extended Data Fig. 9), lending itself a versatile tool for cell and miRNA research. Furthermore, miRNAs regulate various diseases and disorders. RfxCas13d is recently reported to silence neurodegeneration-associated genes in the brain^30^ and alleviate disease-related phenotypes in Huntington’s disease models with minimal off-target effects^25^. Pri-miRNA editing by iREPRESS can inhibit cancer cell proliferation, migration and invasion (Extended Data Fig. 8). As such, dRfxCas13d-based iREPRESS may be exploited to modify pri-miRNAs associated with cancer or other diseases. One challenge for human use is the pre-existing immunity against RfxCad13d^31^, which may mitigate the editing efficacy of iREPRESS in humans. This obstacle may be circumvented by replacing dRfxCad13d with other dCas13 protein capable of RNA targeting such as a small dCas13X.1 (445 aa)^32^. Another roadblock is the collateral effects induced by dRfxCas13d, which is not observed in this study, probably because the fusion of dRfxCas13d with ADAR2_DD_ altered the specificity of dRfxCas13d. To further ease the concern, iREPRESS may be improved by replacing dRfxCas13d with a new high-fidelity Cas13 variant with minimal collateral effects^33^.

## Online content

Any methods, additional references, Nature Research reporting summaries, source data, extended data, supplementary information, acknowledgements, peer review information; details of author contributions and competing interests; and statements of data and code availability are available online.

## Acknowledgements

We would like to thank the Laboratory Animal Center, Chang Gung Memorial Hospital, Linkou, Taiwan for the molecular imaging and technical support. The authors acknowledge the financial support from the National Science and Technology Council (NSTC 112-2223-E-007-002, 111-2223-E-007-003, 111-2634-F-007-007, 111-2622-E-007-003, 109-2314-B-007-002-MY3), Chang Gung Memorial Hospital (CMRPG3I0183, CMRPG3M0781) and National Health Research Institutes (NHRI-EX110-11014BI, NHRI-EX111-11014BI, NHRI-EX112-11014BI), Taiwan.

## Author Contributions

V.A.T designed and performed experiments and wrote the paper. Y.H.C. wrote the paper. Y.T., T.K.N.N., N.N.P., C.W.C., Y.H.C., T.K.D.N., H.D.P., J.T., T.Q.D., A.V.T. performed experiments. Y.C.H. supervised the project and wrote the paper,

## Materials and methods

### Construction of REPRESS plasmids and recombinant BV

The loxP-CMV enhancer-EF1α promoter element^21^, dPspCas13b, human ADAR2_DD_ E488Q (referred to as ADAR2_DD_ in this article) from pC0039 (Addgene #133849)^6^, and WPRE-SV40 poly A signal-loxP^21^ were PCR-amplified with PrimeSTAR Max DNA polymerase (Takara). The N-terminus of dPspCas13b and the C-terminus of ADAR2_DD_ were tethered with the bipartite nuclear localization signal sequence to facilitate nuclear translocation. Amplicons were joined with NEBuilder HiFi DNA assembly master mix (NEB) to the templated REPRESS plasmid (Fig. 1A). Various peptide linkers between dPspCas13b and ADAR2_DD_ was introduced via inverse PCR with divergent primers appended with desired peptide sequence, followed by phosphorylation with T4 Polynucleotide Kinase (NEB). The phosphorylated amplicons were self-ligated with RBC Rapid Ligation kit (RBC Bioscience) at room temperature for 30 min to generate B-REPRESS 1 to 6 plasmids (Fig. 1B). Similarly, D-REPRESS 1 to 6 plasmids were cloned following the same manner with dRfxCas13d amplification from pXR002 (Addgene #109050)^24^. The full-length FL hADAR2 was cloned from pmGFP-ADAR2 (Addgene # 117929)^34^. dEXPRESS plasmid was constructed by replacing ADAR2_DD_ E488Q in pD-REPRESS 5 with deactivated ADAR2_DD_ E488Q E396A via inverse PCR and self-ligation.

Spacer sequences were selected based on two criteria: (i) the relative distance between the target site and the basal junctions of pre-miRNA hairpin and (ii) scores for Cas13d guide design using web-based algorithm (https://cas13design.nygenome.org/^5, 35^). The selected sequences (Supplementary Table 1) were chemically synthesized and cloned into *Bbs*I-digested pC0043 (encoding crRNA for dPsbCas13b) (Addgene #103862) or pXR004 (encoding precursor crRNA for dRfxCas13d) (Addgene #109054) to generate pB-crRNA or pD-crRNA plasmids. Plasmids expressing arRNA and gRNA (for LEAPER^7^ and RESTORE^8^ systems, respectively) were constructed by annealing chemically synthesized oligonucleotides and subcloning into *EcoR*I/*Bbs*I-digested psgRNAa^36^. Alternatively, pXR004 carrying the original *Bbs*I cloning sites was used as a non-targeting (NT) crRNA.

Donor plasmids for recombinant bacmid and BV production were generated by joining two PCR amplicons: one from the entire REPRESS from D-REPRESS 5 or dREPRESS expression cassette; and the other from the donor backbone of pFastBac^®^ Dual expression vector (Gibco). The PCR amplicon joining yielded pBac-REPRESS and pBac-dREPRESS. Similarly, expression cassette harboring the pri-miR-21-targeting (cr21) or non-targeting (NT crRNA) spacers (Supplementary Table 1) were joined with the donor backbone of pFastBac^®^ to generate pBac-cr21 or pBac-NT crRNA.

### Cell culture, transfection and electroporation

HEK293FT (Invitrogen) and A549 cells were maintained in DMEM high glucose (Gibco) supplemented with 10% fetal bovine serum (FBS) and 1% penicillin-streptomycin (PS) (Gibco). Cells were passaged at ≍ 80% confluency and seeded to 12-well plates (1×10^5^ cells/well). After overnight culture, 4 sets of plasmids were co-transfected into cells using Lipofectamine 3000 (Invitrogen) with a DNA:lipid ratio of 1:1.5: (i) REPRESS (300 ng) and crRNA (600 ng) plasmids; (ii) pC0039 (300 ng) and crRNA (600 ng) plasmids for REPAIR; (iii) 500 ng of U6-driven arRNA plasmid for LEAPER; or (iv) pmGFP-ADAR2 (250 ng) and U6-driven gRNA plasmid (750 ng) for RESTORE.

ASCs were isolated from SD rats (Biolasco, Taiwan) as described^37^. The cells were cultured in complete αMEM (Gibco) containing 10% FBS, 1% PS and 4 ng/mL basic fibroblast growth factor (bFGF). ASCs at passage 2 to 5 were used for subsequent experiments. ASCs were resuspended in Opti-MEM (Gibco) and mixed with 3 µg of REPRESS and 6 µg of crRNA plasmids for REPRESS; 3 µg of pC0039 and 6 µg of crRNA plasmid for REPAIR; 5 µg of U6-driven arRNA plasmid for LEAPER; or 2.5 µg of pmGFP-ADAR2 and 7.5 µg of U6-driven gRNA plasmids for RESTORE. The plasmids were electroporated into cells in a 2-mm gap cuvette with NEPA21 Electroporator (NEPAGENE) at 2 pulses of 275 V/2.5 ms with an interval of 50 ms for poring phase and 5 pulses of 20 V/50 ms with an interval of 50 ms.

### Recombinant BV production and ASCs transduction

Recombinant BV were produced using the Bac-to-Bac^TM^ system (Invitrogen) as described^38^. Briefly, *E. coli* DH10Bac (Gibco) was transformed with donor plasmids (pBac-REPRESS, pBac-dREPRESS, pBac-cr21 or pBac-NT crRNA) and plated on Luria plate containing gentamycin (10 µg/mL), kanamycin (50 µg/mL), tetracycline (10 µg/mL), X-gal (200 µg/mL) and IPTG (40 µg/mL) at 37°C overnight. White colonies were selected and further cultured in LB broth containing gentamycin, kanamycin, tetracycline of the same concentration. The recombinant bacmids were purified from the cultured cells for infecting SF9 cells to generate P0 viruses. P0 cells were used to infect SF9 cells for BV amplification. Cre-expressing baculovirus (Bac-Cre) was constructed previously^39^.

For transduction of ASCs, cells were seeded to 12-well plates (5×10^4^ cells/well) and cultured overnight at 37°C under 5% CO_2_. The cells were washed and incubated with BV suspended in surrounding medium (complete culture medium without sodium bicarbonate) at optimal multiplicity of infection (MOI) and mixed on a rocking plate (10 rpm, room temperature). After 6 h, the cells were cultured in fresh medium containing 3 mM sodium butyrate. After 18 h, the sodium butyrate-containing medium was removed, and the cells continued to be cultured in fresh or induction medium for subsequent analysis.

### ASCs differentiation and characterization

Transduced ASC cells were differentiated in adipogenic, chondrogenic or osteogenic induction medium. Half of the media was removed and replaced by fresh induction media until analysis time. Differentiated cells were fixed in 4% buffered formaldehyde (Macron) for 15 min at room temperature and stained with Oil Red O, Alcian Blue, or Alizarin Red for adipogenesis, chondrogenesis, or osteogenesis, respectively. The dyes in the stained cells were extracted with isopropanol, guanidine hydrochloride, or cetylpyridinium chloride and read at 492, 595, or 550 nm on the plate reader (SpectraMax M2, Molecular Devices) for quantitation.

### Cell metabolism and proliferation assay

ASCs cells were seeded to 96-well plates (2.5×10^3^ cells/well) prior to transduction. Mock or transduced cells were cultured in fresh medium and replenished every 2 days. For metabolism assay, cells were washed with PBS and incubated in fresh medium with 0.5 mg/mL MTT (Sigma) at 37°C for 3 h. After medium removal, cells were incubated with 100 µL solution of 4 µM HCl and 0.1% Triton X-100 in isopropyl alcohol at room temperature for 15 min on a rocking plate. The adsorption was read on SpectraMax M2 at 560 nm. Alternatively, cells were stained with BrdU for proliferation assay (Cell BioLabs). Briefly, the stained cells were fixed and subsequently stained with anti-BrdU antibody followed by HRP-conjugated secondary antibody staining. The cells were allowed to react with substrate. The reaction was stopped before reading at 450 nm on SpectraMax M2.

### Adenine conversion rate by Sanger sequencing

Total RNA was extracted with Gene-Spin™ Total RNA Purification Kit (Protech). cDNA synthesis was performed using pri-miRNA-specific primers (Supplementary Table 2) with High-Capacity cDNA Reverse Transcription Kit (Applied Biosystems) to amplify the pre-miRNA region of interest using the following temperature cycle: RNA and the primer mixtures were heated at 65°C/5 mins and incubated at 4°C prior to the addition of reaction mixture, followed by 37°C/2 h and 85°C/2 min. The cDNA was PCR-amplified with primers flanking the pre-miRNA region (Supplementary Table 2). The amplicons were purified, and Sanger sequenced (Genomics, Taiwan). The ab1. chromatogram files were used to calculated A conversion rate by EditR^40^.

### Quantitative reverse-transcription PCR

cDNA was synthesized from total RNA using random hexamers with High-Capacity cDNA Reverse Transcription Kit (Applied Biosystems). Gene expression quantitation was performed with qPCRBIO Probe Blue Mix Lo-ROX (PCRBIOSYTEMS) on LightCycler^®^ 96 (Roche) with primers specific for *C/ebpα*, *Ppar-γ*, *Acan*, *Col2a1*, *HOX3B* and *Vmp1*.

Alternatively, miRNA-enriched total RNA was collected using NucleoSpin miRNA Plus Mini Extraction Kit (MACHEREY NAGEL). miRNA amplification and expression were performed with TaqMan miRNA assay (Thermo) as instructed and measured on LightCycler^®^ 96 (Roche).

### Amplicon library preparation and analysis

The cDNA reverse transcribed from total RNA and PCR amplified with pre-miRNA-flanking primers carrying TruSeq indexes for high throughput sequencing. PCR amplification was carried out in 25-µL reaction of PrimeSTAR Max DNA polymerase (Takara), containing 0.2 µM of forward and reverse primers. 5 µL of each amplicon were resolved on 2% agarose gel and pooled together in equivalent amount prior to purification by GeneJET PCR Purification Kit (Thermo). Library preparation was performed with TruSeq® Nano DNA Library Prep (Illumina) (Genomics, BioSci & Tech Co, Taiwan). Library quality was assessed on a Qubit 2.0 Fluorometer (Thermo Scientific) and Agilent Bioanalyzer 2100 and sequenced on NovaSeq 6000 (Illumina) with 150 bp paired-end reading. Paired-end reads were trimmed by Trimmomatics^41^ and demultiplexed by in-house script. Demultiplexed paired-end reads were joined with PEAR v0.9.6^42^, and unique joined sequences were counted by in-house Perl script.

### Transcriptome sequencing and analysis

Total RNA was used for library preparation by TruSeq Stranded mRNA Library Prep Kit (Illumina). Briefly, mRNA was purified from 1 μg of total RNA by oligo(dT)-coupled magnetic beads and fragmented into small pieces under elevated temperature. cDNA was synthesized using reverse transcriptase and random primers, followed by double stranded cDNA generation, 3′ adenylation and adapter ligation. The library was purified with AMPure XP system (Beckman Coulter) and the quality was assessed on Qsep400 with N1 High Sensitivity Cartridge Kit (Bioptic Inc., Taiwan). The qualified library was sequenced on NovaSeq 6000 (Illumina) with 150 bp paired-end reading (Genomics, BioSci & Tech Co., Taiwan). Bases with low quality and sequences from adapters in raw data were removed using program fastp (version 0.20.0)^43^. The filtered reads were aligned to the reference genomes using HISAT2 (version 2.1.0)^44^. FeatureCounts (v2.0.1) in Subread package was applied for the quantification of gene abundance^45^. EdgeR was used to perform differential gene expression (DGE) analysis^46^.

### Small RNA sequencing and analysis

miRNA-enriched total RNA (100 ng) was used for small RNA library preparation with QIAseq® miRNA Library Kit (QIAGEN) as instructed. Briefly, 3′ and 5′ adapters were directly and specifically ligated to 3′ and 5′ end of small RNA, respectively. First-strand cDNA was synthesized using QIAseq miRNA NGS RT Enzyme and Primer. cDNA was purified by QIAseq beads prior to PCR amplification with QIAseq miRNA NGS Library Buffer and HotStarTaq^®^ DNA Polymerase. The library was purified by QMN Beads and the quality was assessed on the Qsep400. The qualified library was sequenced on NovaSeq 6000 (Illumina) with trimmed 75 bp single-end reading (Genomics, BioSci & Tech Co.). Adapter sequences were trimmed with TrimGalore! And mapped to miRBase reference with bowtie to obtain proper miRNA reads^47–49^. Sam/bam files were processed with SAMtools and the expression profile of miRNAs are calculated and normalized with EdgeR^46, 50^. Differentially expressed miRNAs (DEmiRNAs) are identified using DEGSeq^51^.

### Cell/scaffold construct preparation and characterization

ASCs were mock or co-transduced in a 15-cm dish as described previously. At 1 dpt, gelatin discs (d=6 mm, thickness 1 mm) were cut from Spongostan sheets (Ethicon, MS0003) with a biopsy puncher (Integra Miltex), and prewetted in PBS. ASCs cells were collected and seeded to gelatin discs (5×10^6^ cells/disc). The resultant ASCs/gelatin constrcts were incubated at 37°C, 5% CO_2_ for 4 h prior to culturing in chondroinduction medium. At 7 and 14 dpt, the constructs were collected for calvarial defect implantation. Alternatively, the constructs were fixed in 4% buffered formaldehyde (Macron) overnight at room temperature before embedding in paraffin and sliced into 3-µm thick sections. The sections were cleaned in xylene and hydrated through a series of descending alcohol. For histochemical analysis, the hydrated sections were stained with H&E and Alcian Blue/Nuclear Fast Red. Alternatively, antigen retrieval of the sections were carried out with 0.05% trypsin EDTA (Gibco) at 37°C for 20 min followed by blocking in PBS containing 5% bovine serum albumin and 0.1% Tween 20 for 1 h before incubating with rabbit anti-collagen type II (Abcam, 1:100) primary antibody at 4°C overnight. The sections were then incubated with HRP-conjugated goat anti-rabbit (Abcam, 1:500) secondary antibody at room temperature for 1 h and developed with SIGMAFAST™ 3,3′-Diaminobenzidine (Sigma). Sections were photographed on Eclipse TS100-F (Nikon). Positively stained areas of the images were processed by Fiji for semi-quantification.

### Animal implantation

Animal experiments were performed in compliance with the Guide for the Care and Use of Laboratory Animals (National Council of Science and Technology, Taiwan) and were approved by the Institutional Animal Care and Use Committee of National Tsing Hua University. Mock (Mock group) and transduced (iREPRESS) ASCs/gelatin constructs cultured in chondroinduction medium were harvested at 7 or 14 dpt for implantation. On the day of implantation, 6-week-old SD rats were injected intramuscularly with Zoletil^®^ 50 (25 mg/kg body weight, Virbac Animal Health) and 2% Rompun^®^ (0.15 mL/kg body weight, Bayer Health Care) for anesthesia, followed by cefazolin injection (160 mg/kg body weight). The calvaria were exposed by a 2-cm midline incision and the removal of the periosteal layers.

A 6-mm defect was created by punching through the exposed calvarium with a biopsy puncher (Integra Miltex) gradually with occasional saline buffer supplement to minimize damage to the neighboring bone tissue and underlying dura mater. The bone flaps were gently discarded and replaced with the ASCs/scaffold constructs followed by suturing with 4-0 absorbable stitch (PolySorb, Covidien). Post-operative procedure was performed with an administration of topical neomycin/bacitracin and intramuscular injection of ketoprofen (5 mg/kg).

### Qualitative and quantitative characterization of bone healing

µCT imaging was performed on a nanoScan SPECT/CT (Mediso) at Chang Gung Memorial Hospital to assess bone regeneration of the animals at 4 (W4), 8 (W8) and 12 (W12) weeks post-implantation. 3D and hot-and-cold projections of the calvarial bones were rendered by InterView^TM^ FUSION (Mediso) and PMOD (PMOD Technologies). DICOM files from scanning were further processed by PMOD to generate bone area, volume and density.

### Statistical analysis

*In vitro* data and images are representative of at least three independent culture experiments. All quantitative data are expressed as mean±standard deviation (SD). Statistical significance was evaluated by Student’s *t* test, one-way ANOVA, or two-way ANOVA analyses. A *p* value less than 0.05 was considered significant.

## Extended Data

**Extended Data Fig. 1.**
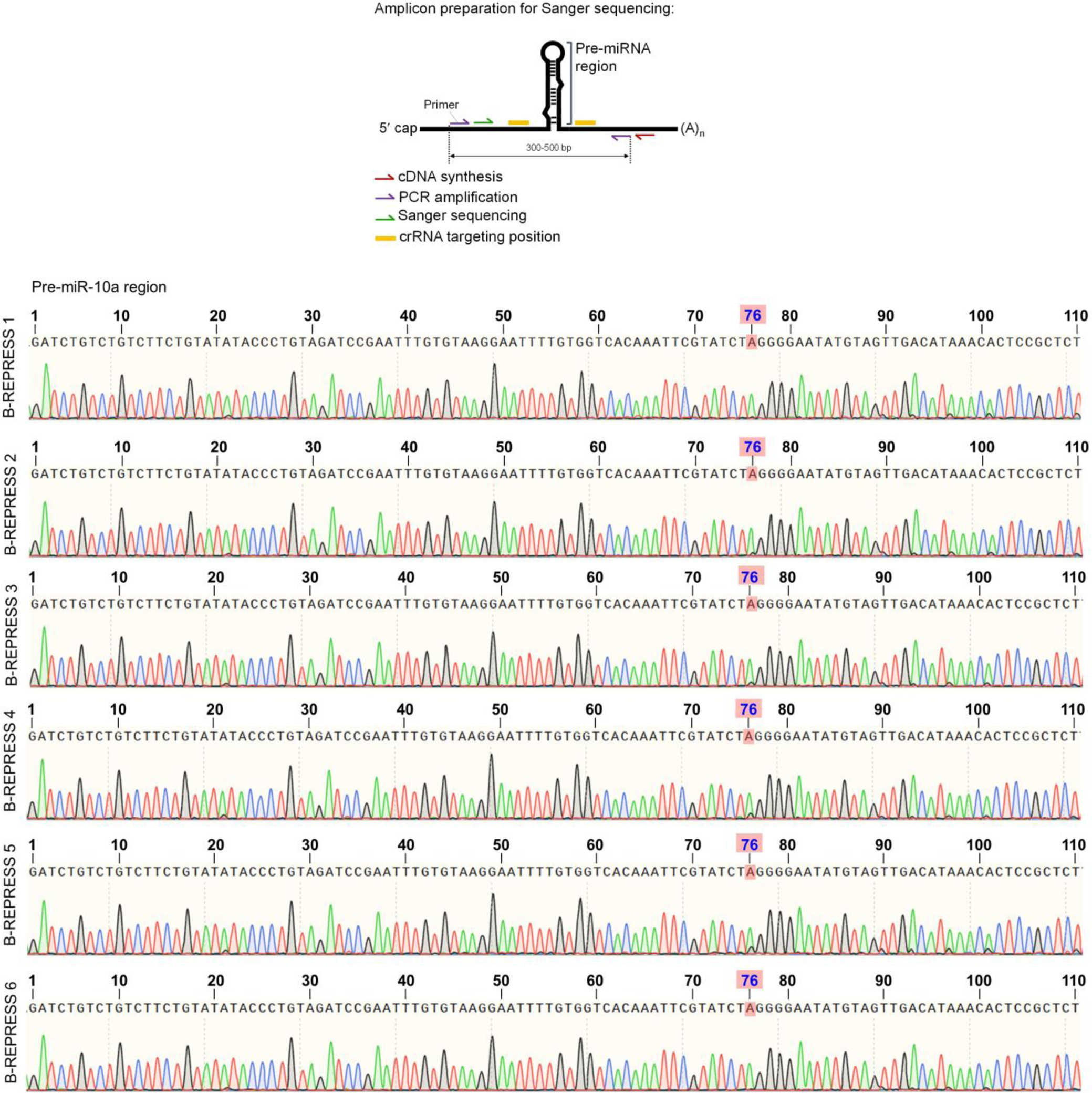
A->I conversion by B-REPRESS in pri-miRNA-10a. HEK293FT cells were co-transfected with pB-REPRESS and pB-crRNA plasmids. After 1 day, total RNA was reverse transcribed with pri-miR-10a-specific primer (primer for cDNA synthesis). The cDNA encompassing the pre-miRNA and 5’ and 3’ flanking sequences was PCR-amplified using primer pairs flanking the pre-miR-10a hairpin region (primers for PCR amplification). PCR amplicons (444 bp) were sequenced using primers for Sanger sequencing and the .ab1 files were uploaded to https://moriaritylab.shinyapps.io/editr_v10/ for editing efficiency analysis. For simplicity, representative chromatograms only show the pre-miR-10a region sequence while other sequences are not shown. The data show A->I editing events only at A_76_ (two peaks). No other edits were observed outside the pre-miR-10a region, including the crRNA targeting positions.

**Extended Data Fig. 2.**
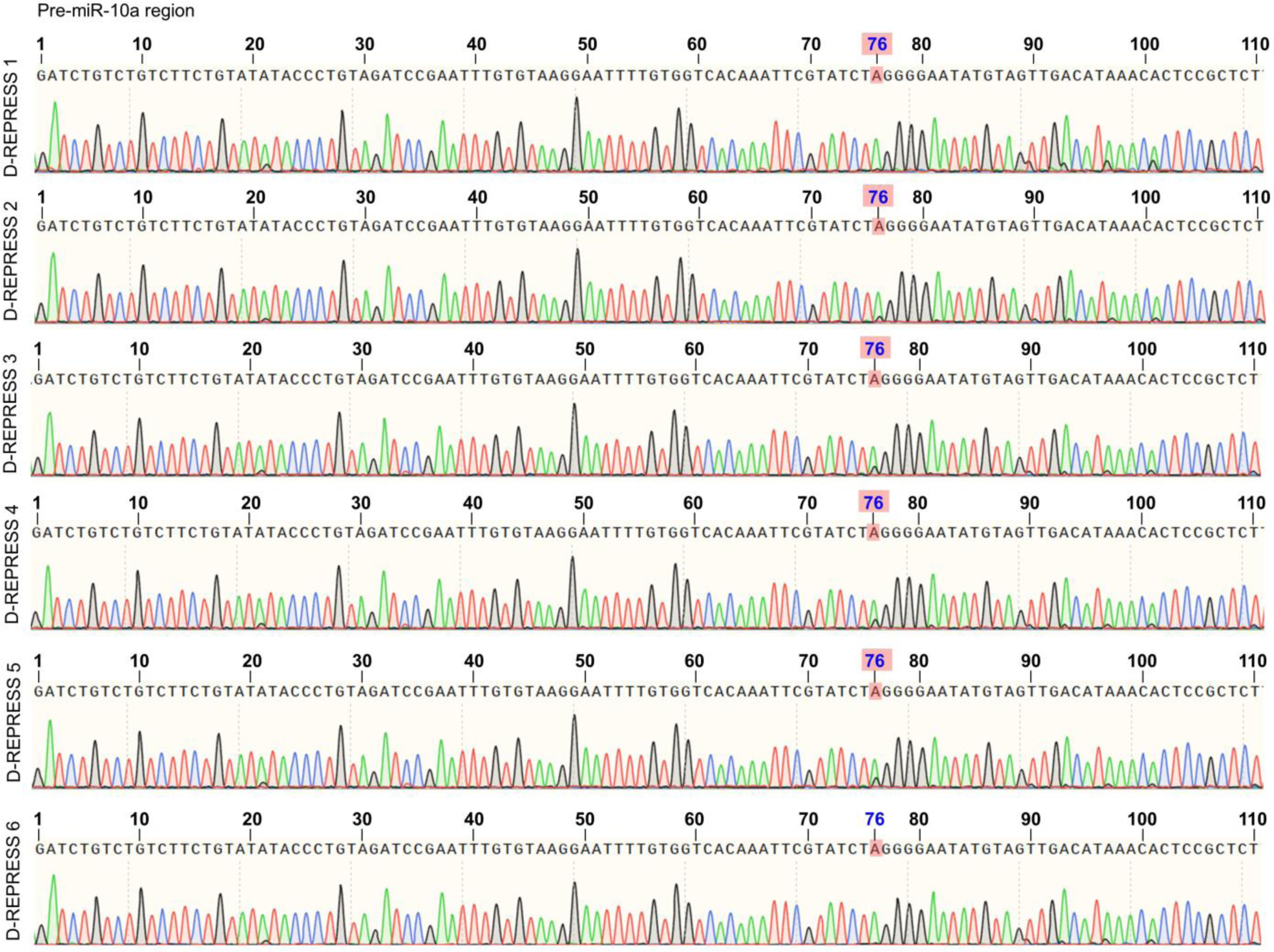
A->I conversion by D-REPRESS in pri-miRNA-10a. HEK293FT cells were co-transfected with pD-REPRESS and pD-crRNA plasmids. Subsequent procedures for cDNA synthesis, PCR amplification and Sanger sequencing were identical to those in Extended Data Fig. 1. For simplicity, representative chromatograms only show the pre-miR-10a region sequence while other sequences are not shown. The data show A->I editing events only at A_76_ (two peaks). No other edits were observed outside the pre-miR-10a region, including the crRNA targeting positions.

**Extended Data Fig. 3.**
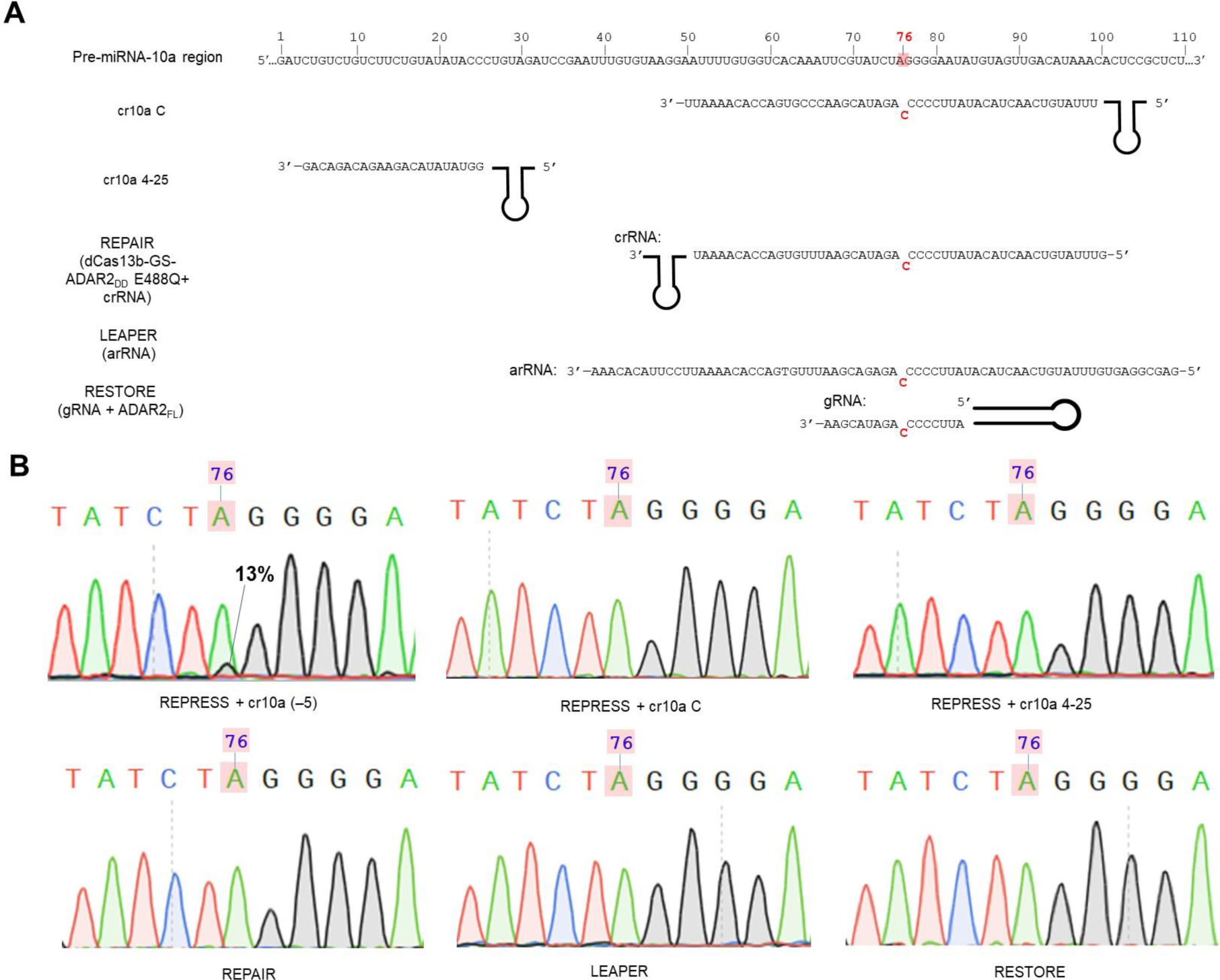
A->I conversion by direct targeting at the pre-miR-10a hairpin. This experiment aimed to test whether RNA guide to target the pre-miR-10a hairpin enables A->I conversion. (A) Illustration of targeting sites and RNA guide designs for REPRESS, REPAIR, LEAPER and RESTORE. Cr10a C and cr10a 3-25 are crRNAs separately co-transfected with pD-REPRESS 5 into HEK293FT cells. cr10a C encodes a 50-nt spacer with a C mismatch opposite to A_76_ of the pre-miR-10a. cr10a 3-25 encodes a 22-nt spacer with the highest targeting score (0.88) and fully complementary to nucleotides 3 to 25 of pre-miR-10a. The remaining RNA sequences were designed following their respective original systems. For REPAIR, dPspCas13b-GS-ADAR2_DD_ and crRNA plasmids were co-transfected into HEK293FT. For LEAPER, U6-driven arRNA expression plasmid was transfected. For RESTORE, U6-driven gRNA and CMV-driven full length ADAR2 (ADAR2_FL_) expression plasmids were co-transfected. PCR amplicons were generated as described in Extended Data Fig. 1 for Sanger sequencing. (B) Sanger sequencing chromatograms show that A->I conversion occurs at A_76_ only by REPRESS with a crRNA targeting ssRNA motif (*d*=−5, as shown in Fig. 1H). No A->I conversion higher than background levels was detected for REPRESS using Cr10a C and cr10a 3-25, REPAIR, LEAPER and RESTORE. Translucent red box indicates the intended edited adenine.

**Extended Data Fig. 4:**
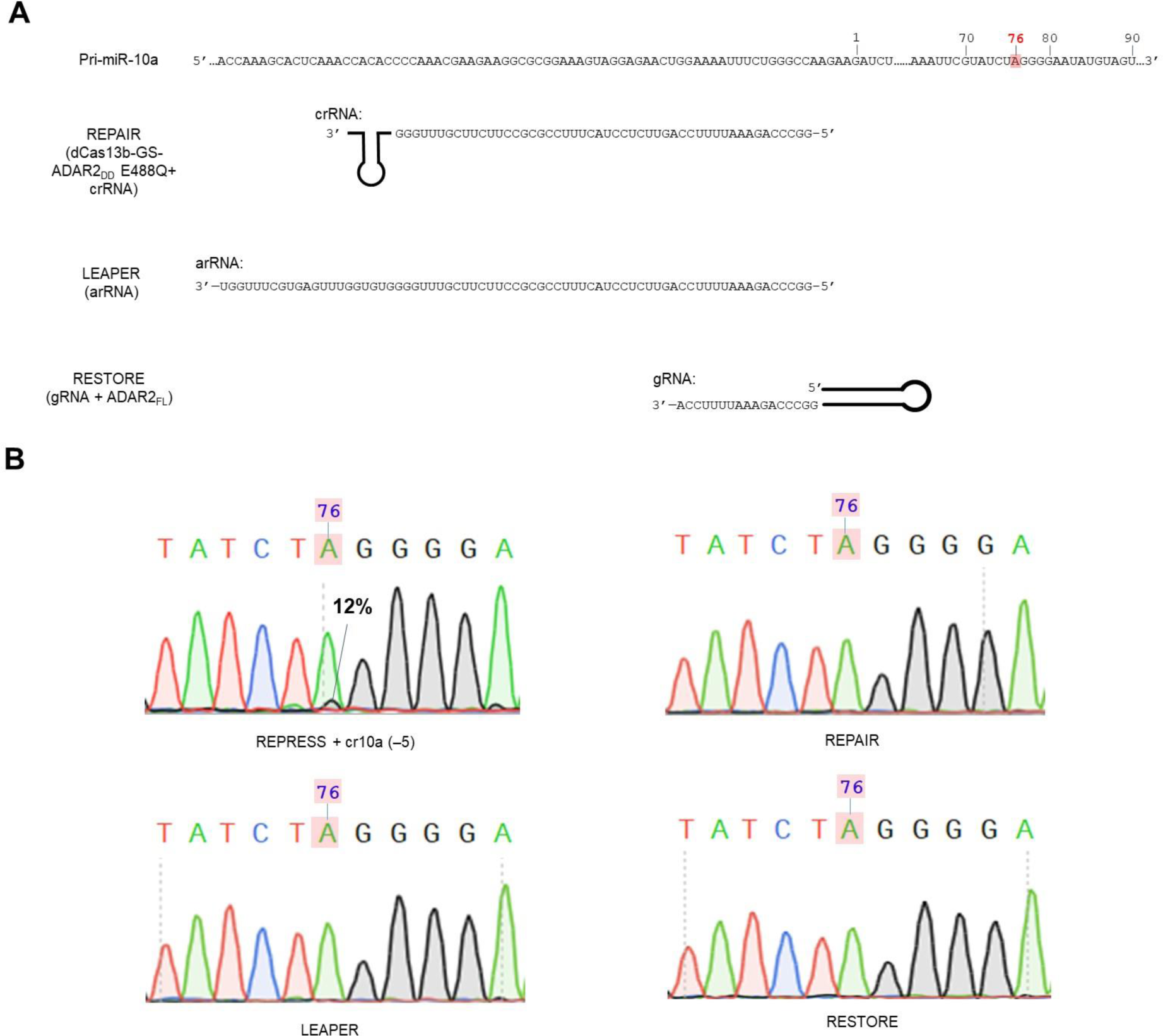
A->I editing with proximal targeting near the pre-miR-10a hairpin. This experiment aimed to test whether RNA guide to target the ssRNA sequences in the vicinity of basal junction of pre-miR-10a hairpin enables A->I conversion. (A) Illustration of targeting sites and RNA guide designs for REPAIR, LEAPER and RESTORE. These RNA guides were designed to mimic the REPRESS design and target the ssRNA close to the 5’ basal junction of pre-miR-10a. The length of each RNA was selected following its respective original system. (B) Sanger sequencing chromatograms show editing results of REPRESS, REPAIR, LEAPER, and RESTORE. Experimental procedures were identical to those in Extended Data Fig. 3. For REPAIR, dCas13b-GS-ADAR2_DD_ and crRNA plasmids were co-transfected into HEK293FT. For LEAPER, U6-driven arRNA expression plasmid was transfected. For RESTORE, U6-driven gRNA and CMV-driven full length ADAR2 (ADAR2_FL_) expression plasmids were co-transfected. PCR amplicons were generated as described as in Extended Data Fig. 1 for Sanger sequencing. Translucent red box indicates the intended edited adenosine. Only REPRESS enabled A->I conversion at A_76_, while REPAIR, LEAPER, and RESTORE systems failed to achieve A->I editing.

**Extended Data Fig. 5:**
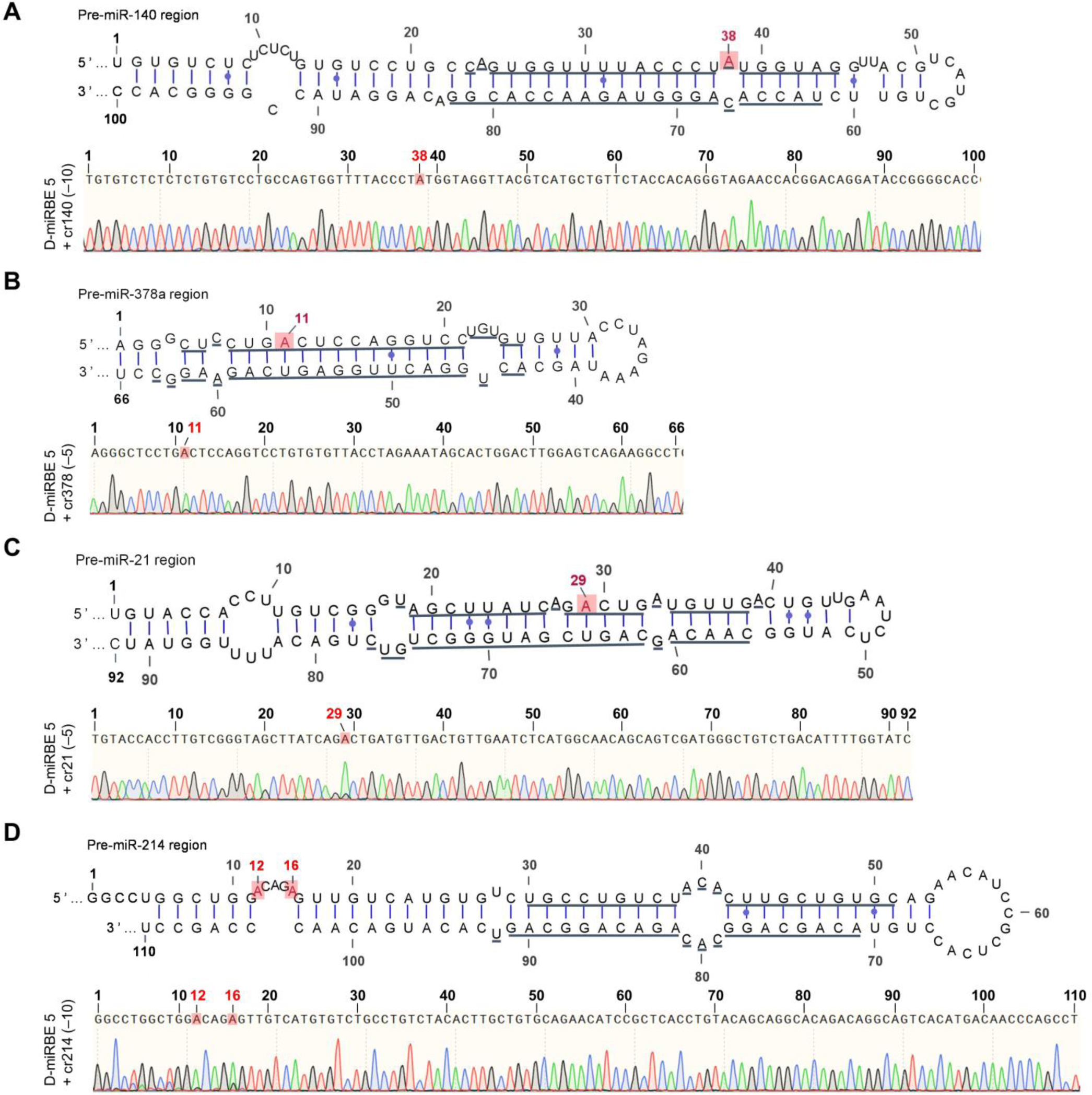
REPRESS enabled editing of differentt pri-miRNAs. The secondary structures of the pre-miRNA stem loops were retrived from https://www.mirbase.org/ and rendered by VARNA. Adenine selection of REPRESS for deamination varied across different pri-miRNAs. While deamination occurs to A within the mature sequence for pre-miR-140 (A_38_), −378a (A_11_), and −21 (A_29_), REPRESS converted A outside the mature sequence of pre-miR-214 (A_12_/A_16_). The adenine deamination may occur at A with a C mismatch (pre-miR-140, A_38_-C) or with paired A·U (pre-miR-378a and −21). The A->I conversion also occurred at A in the bulge loop (pre-miR-214), which was also observed previously (Yang*, et al.*, 2006).

**Extended Data Fig. 6.**
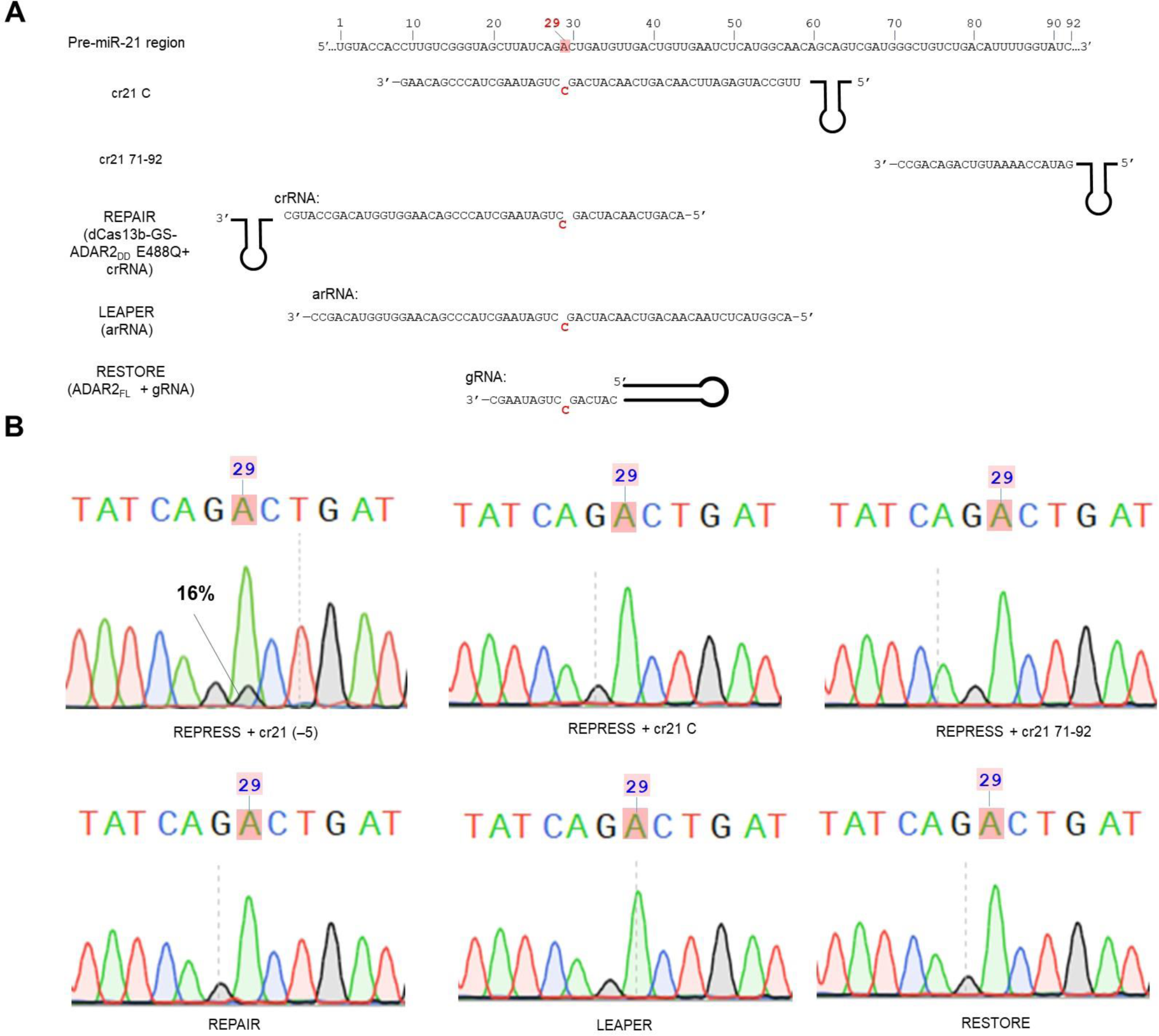
A->I conversion of pri-miR-21 in ASCs by direct targeting. This experiment aimed to explore whether RNA guide to target the pre-miR-21 hairpin enables A->I conversion. (A) Illustration of targeting sites and RNA guide designs for REPRESS, REPAIR, LEAPER and RESTORE. Cr21 C encodes a 50 nt spacer carrying a C mismatch opposite A_29_ of pre-miR-21. Cr21 71-92 encodes a 22 nt spacer with the highest targeting score (1.0) and fully complementary to position 71 to 92 of pre-miR-21. The remaining RNA sequences were designed following their respective original systems. For REPAIR, dPspCas13b-GS-ADAR2_DD_(E488Q) and crRNA plasmids were co-electroporated into ASCs. For LEAPER, U6-driven arRNA expression plasmid was transfected. For RESTORE, U6-driven gRNA and CMV-driven full length ADAR2 (ADAR2_FL_) expression plasmids were co-transfected. PCR amplicons were generated as described above for Sanger sequencing. (B) Sanger sequencing chromatograms show that A_29_ conversion only occurs when using REPRESS with cr21 (−5) that targets the ssRNA sequence near the basal junction (*d*=−5). However, REPRESS, REPAIR, LEAPER and RESTORE directly targeting the pre-miR-21 hairpin fail to catalyze A_29_ conversion. Translucent red box indicates the intended edited A.

**Extended Data Fig. 7.**
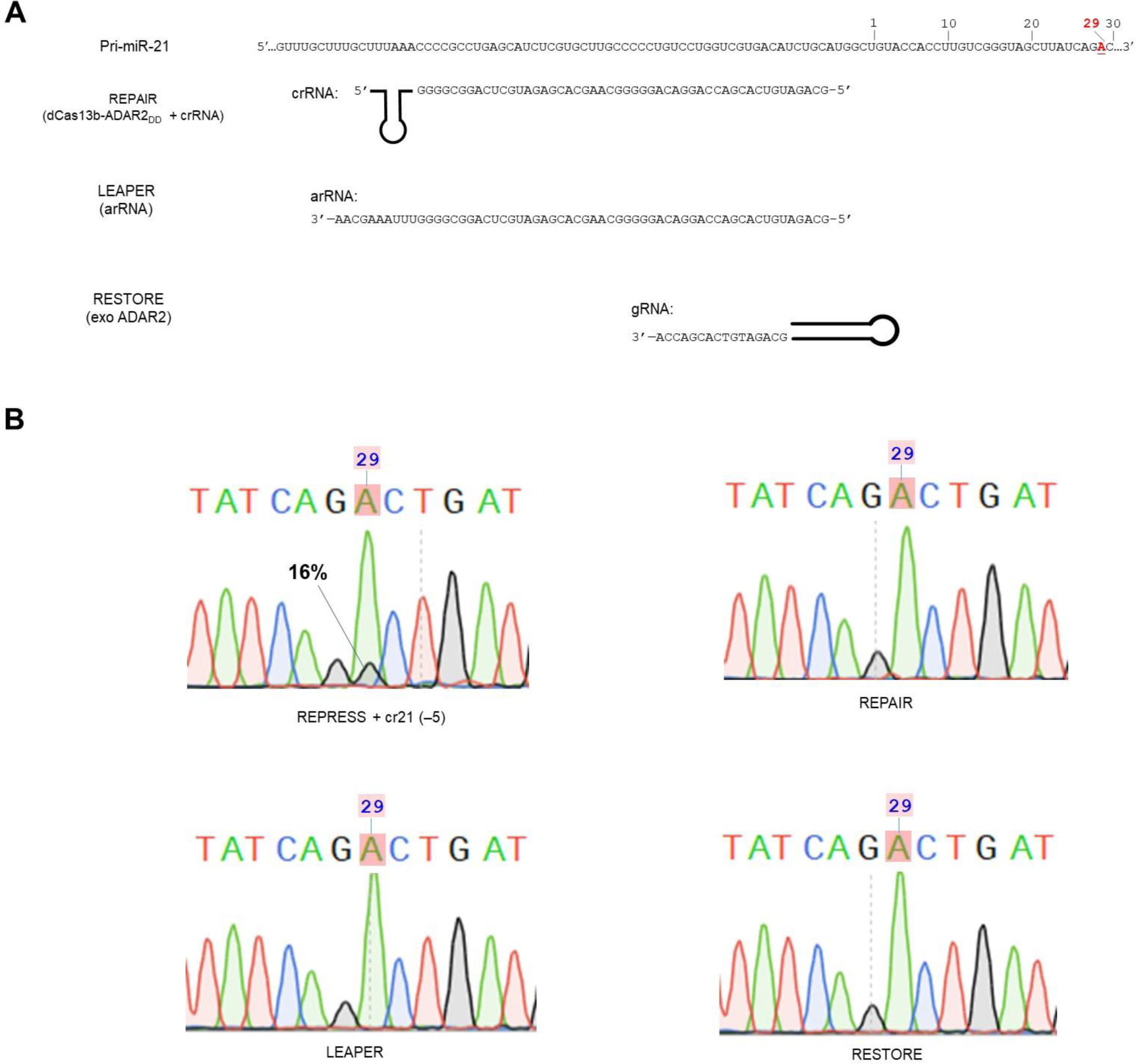
A->I conversion of pri-miR-21 in ASCs by proximal targeting. This experiment aimed to explore whether RNA guide to target the ssRNA sequence near the basal junction of pre-miR-21 hairpin by other systems enables A->I conversion. (A) Schematic illustration of the targeting site and RNA guide designs for REPAIR, LEAPER, and RESTORE. The length of each RNA was selected following its respective original system. ASCs were electroporated with plasmids for these experiments. The experiments were performed as in Extended Data Fig. 6. (B) Sanger sequencing chromatograms show that A_29_ conversion only occurs when using REPRESS with cr21 (−5) that targets the ssRNA sequence near the basal junction (*d*=−5). However, REPRESS, REPAIR, LEAPER and RESTORE directly targeting the pre-miR-21 hairpin fail to catalyze A_29_ conversion. Translucent red box indicates the intended edited adenosine.

**Extended Data Fig. 8:**
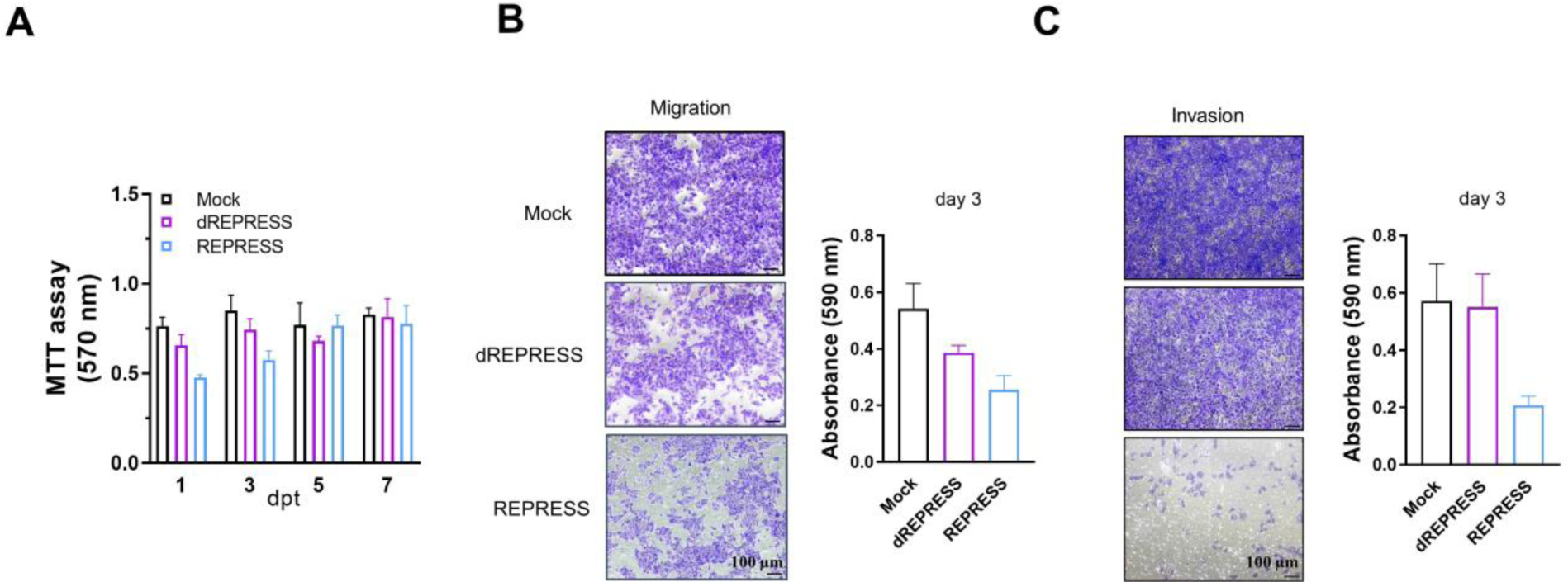
REPRESS-mediated miR-10a knockdown inhibited lung cancer cell A549 proliferation, migration and invasion. A549 was transfected with plasmids encoding REPRESS or dREPRESS (300 ng) and crRNA targeting pri-miR-10a (600 ng). (A) Proliferation assay of transfected cells. One day after transfection, cell proliferation in the REPRESS group was inhibited as compared to Mock or dREPRESS groups, but began to proliferate again at days 3, 5 and 7. (B) Migration assay. (C) Invasion assays. One day after transfection, mock and transfected cells were trypsinized, seeded onto a transwell insert without (B) or with (C) Matrigel, and cultured in complete culture medium. At day 3, cells in the transwell were fixed. The fixed cells in the upper surface of the transwell were scraped away and the lower surface was stained with Crystal Violet for imaging and quantitation. The data show that REPRESS inhibited A549 cells proliferation temporarily and repressed migration and invasion.

**Extended Data Fig. 9.**
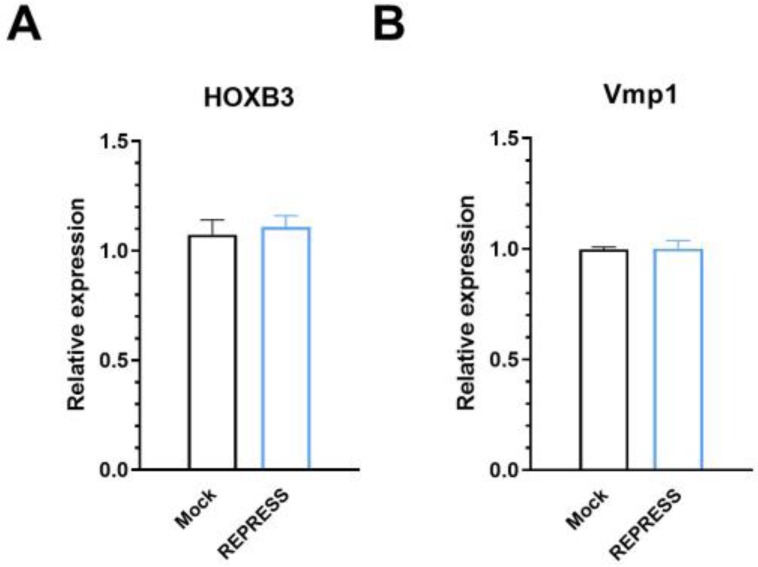
Editing of pri-miRNA did not alter the levels of their host genes. Pri-miRNAs are usually co-expressed with their host genes. HOXB3 and Vmp1 are host genes of miR-10a and miR-21, respectively. (A) Expression levels of HOXB3 in HEK293FT. HEK293FT cells in a 12-well plate were transfected with 300 ng of REPRESS and 600 ng of cr10a (‒5) plasmids. (B) Expression levels of Vmp1 in ASCs. ASC cells (1×10^6^ cells) were electroporated with 3 µg of REPRESS and 6 µg of cr21 (‒5). After 1 day, total RNA was harvested and reverse transcribed with random hexamers. The cDNA was subjected to qRT-PCR with qPCRBIO Probe Blue Mix Lo-ROX (PCRBIOSYTEMS) on LightCycler^®^ 96 (Roche). The gene expression levels were normalized to those of Mock groups. The data show that REPRESS catalyzes local A->I conversion at pri-miR-10a and pri-miR-21 (Fig. 1) without disturbing the expression of host genes.

**Extended Data Fig. 10.**
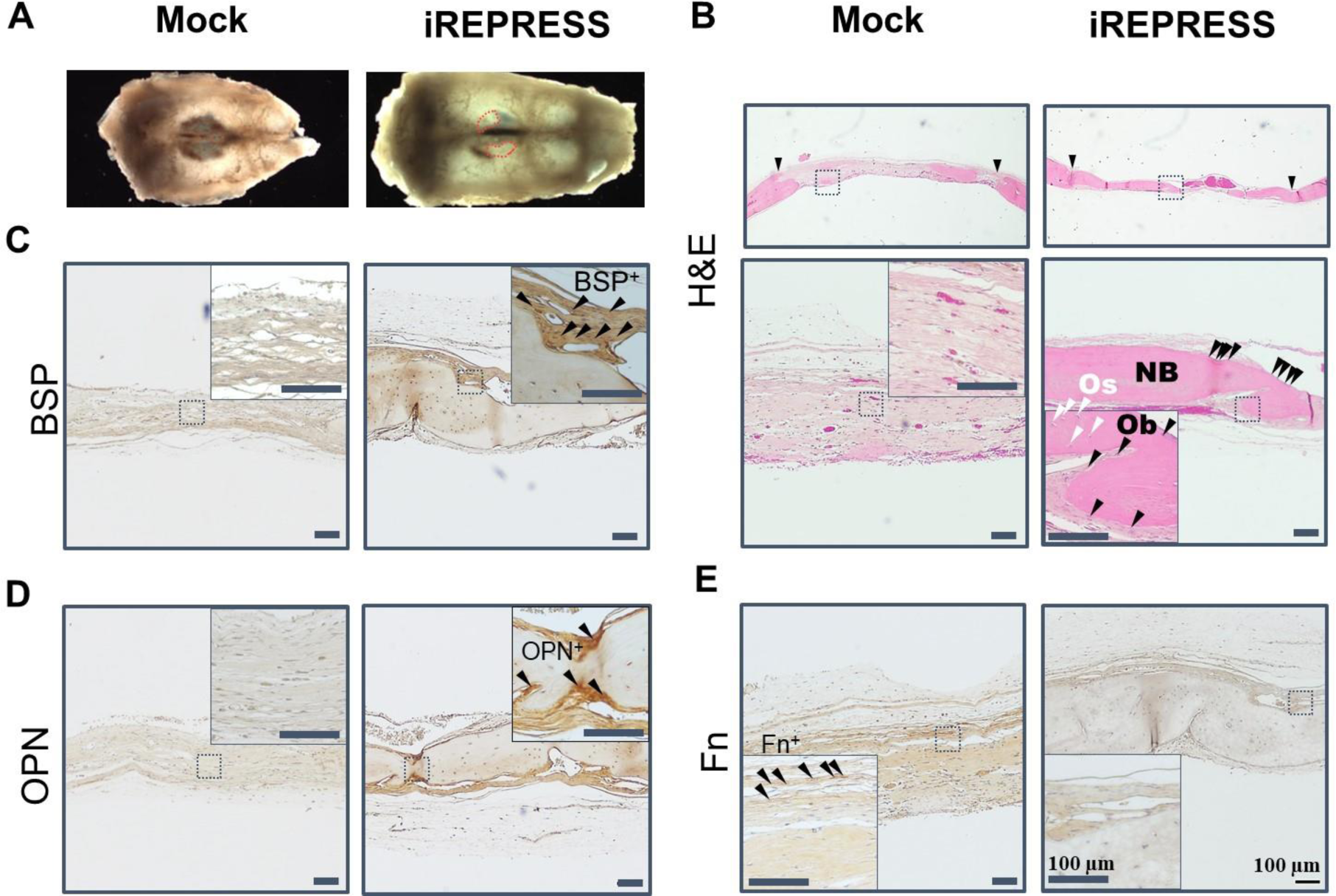
Histochemical and immunohistochemical analyses of bone regeneration. The data show that iREPRESS-engineered cells improved the bone formation, accumulation of bone-specific matrix sialoprotein (BSP) and osteopontin (OPN) and repressed the formation of fibrous tissues. (A) Gross appearance of representative calvarial bones. (B) H&E staining. (C-D) Immunostaining for bone formation markers BSP (C) and OPN. (D) Immunostaining of fibrous tissue marker fibronectin (Fn). Animal experiments were performed as in Fig. 6. Calvarial bones were harvested at W12, fixed in 4% formaldehyde for 2 days, decalcified in Osteosoft (Sigma) for 7 days prior to paraffin embedding and sectioning. The sections were cleaned in xylene and hydrated through a series of descending alcohol. Antigen retrieval was performed with 0.05% trypsin EDTA at 37°C for 20 min followed by blocking in PBS containing 5% bovine serum albumin and 0.1% Tween 20 for 1 h before incubating with rabbit anti-BSP (Abcam, 1:100), rabbit anti-OPN (Abcam, 1:100) or rabbit anti-fibronectin (Abcam, 1:100) primary antibodies at 4°C overnight. The sections were then incubated with HRP-conjugated goat anti-rabbit (Abcam, 1:500) secondary antibodies at room temperature for 1 h prior to developing. The data are representative of 4∼6 bone sections. NB, new bone; Os, osteocyte (white arrowhead); Ob, osteoblast (black arrowhead).

## Supplementary Tables

**Table S1.**
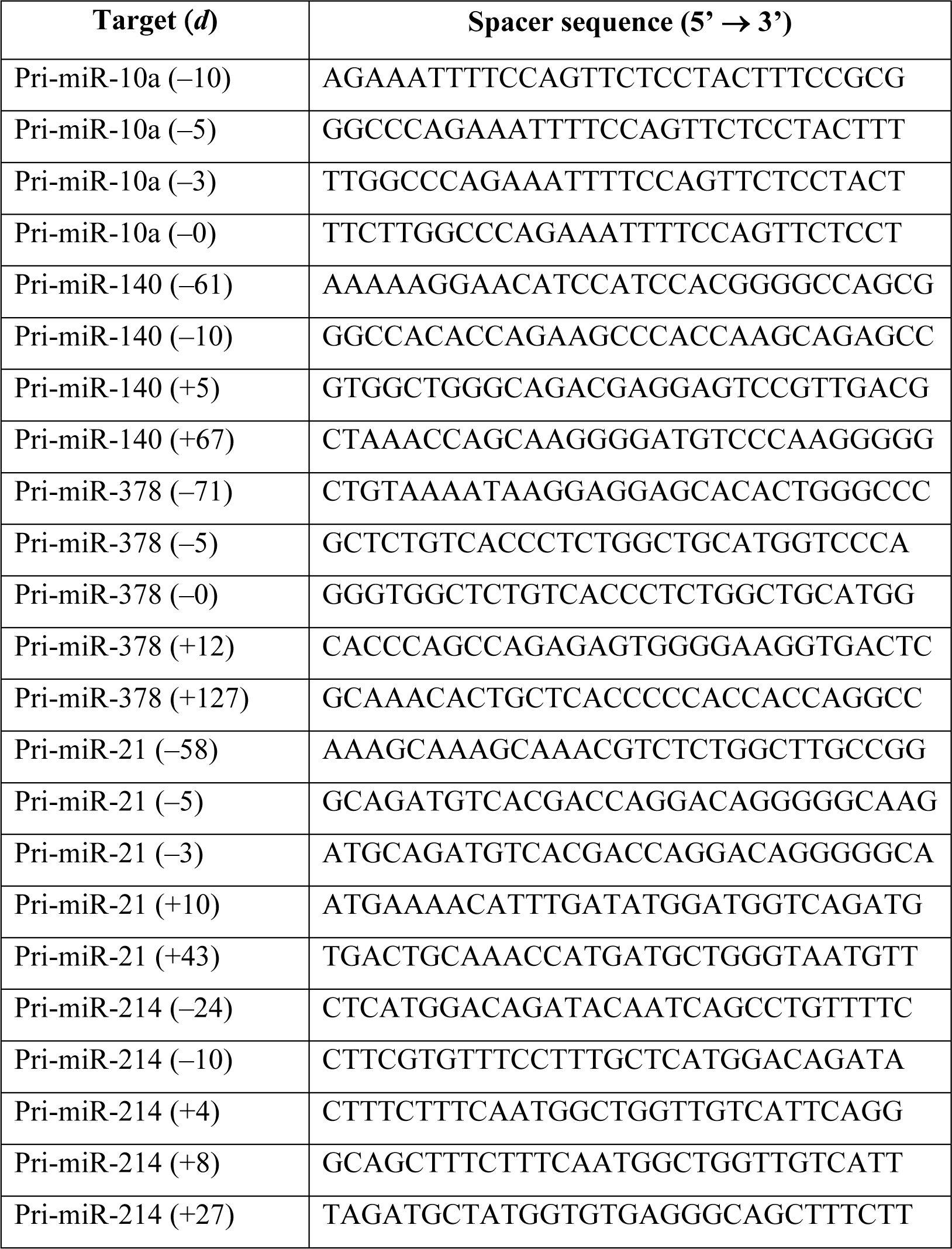
Spacer sequences of the crRNA used in this study.

**Table S2:**
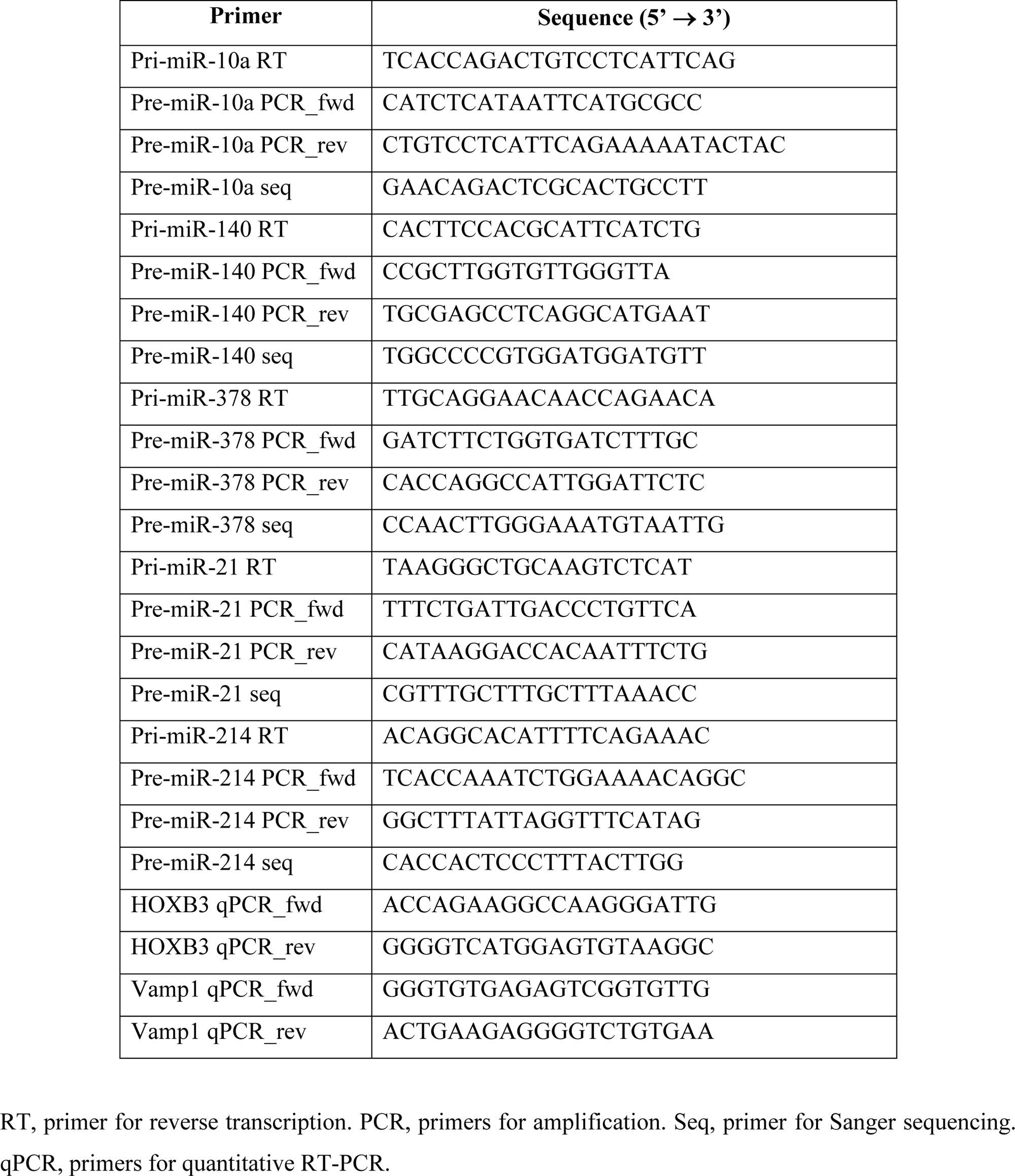
Primer sequences used in this study.

